# Cooperation and deception through stigmergic interactions in human groups

**DOI:** 10.1101/2023.04.20.537511

**Authors:** Thomas Bassanetti, Stéphane Cezera, Maxime Delacroix, Ramón Escobedo, Adrien Blanchet, Clément Sire, Guy Theraulaz

## Abstract

Stigmergy is a generic coordination mechanism widely used by animal societies, in which traces left by individuals in a medium guide and stimulate their subsequent actions. In humans, new forms of stigmergic processes have emerged through the development of online services that extensively use the digital traces left by their users. Here we combine interactive experiments with faithful data-based modeling to investigate how groups of individuals exploit a simple rating system and the resulting traces in an information search task in competitive or non-competitive conditions. We find that stigmergic interactions can help groups to collectively find the cells with the highest values in a table of hidden numbers. We show that individuals can be classified into three behavioral profiles that differ in their degree of cooperation. Moreover, the competitive situation prompts individuals to give deceptive ratings and reinforces the weight of private information versus social information in their decisions.

## Introduction

The exchange of social information is the core mechanism by which groups of individuals are able to coordinate their activities and collectively solve problems [1, 2, 3, 4, 5]. Social information allows individuals to adapt to their environment faster and/or better than through collecting personal information alone [6, 7, 8, 9, 10]. The use of social information thus provides evolutionary advantages to animal groups and occurs in many contexts, such as foraging, decision-making, division of labor, nest building, or colony defense [1, 2, 11, 12].

Quite often, social information is indirectly shared between individuals: some of them leave traces of their activities in the environment and others can use this information to guide their own behavior and inform their own decisions [13]. This form of indirect communication, also called stigmergy, in which the trace of an action left on a medium stimulates the performance of a subsequent action which produces another trace and so on, is widely used by animal societies and especially social insects to self-organize their collective behaviors [14, 15, 16]. These stigmergic interactions that allow the emergence of coordinated activities out of local independent actions should have played a major role in the evolution of cooperativity within groups of organisms [17, 18].

In humans, with the digitalization of society and economies, social information has increasingly taken the form of digital traces, which are the data individuals leave either actively or passively when using the Internet [19, 20, 21]. New forms of stigmergic processes have been identified since these digital traces are largely exploited in social networks and in electronic commerce, in particular through the use of rating and recommendation systems that can help participants to discover new options and make better choices [22, 23, 24, 25, 26]. However, individuals do not use social information in the same way. Some individuals exploit it to make their choices, while others may simply ignore it and only use their own private information, or can even go against the message delivered by social information [27]. In fact, the same individual can even change the way he/she provides and uses social information depending on the context [28].

Moreover, the use of digital traces is very sensitive to noise and manipulation [29, 30]. Indeed, in competitive situations, malicious spammers can manipulate social information by deliberately giving high (respectively, low) ratings to certain low (respectively, high) quality items. Therefore, knowing the way individuals share and use digital traces in different contexts is a crucial step to understanding how groups of individuals can cooperate through stigmergic interactions and can exhibit collective intelligence.

The aims of this study are twofold. First, we study through a combination of experiments and computational modeling how indirect interactions between individuals in a human group involving the use of traces allow them to cooperate during an information search task. Secondly, we study how a competitive or non-competitive context influences the way in which individuals exchange and use the social information embedded in the traces of their past actions to perform the information search task.

Through the development of an interactive web application and the use of data-based modeling, we identify the behavioral and cognitive strategies combined with stigmergic interactions that govern individual decisions. The simulation results of our faithful computational model provide clear evidence that the collective behavioral dynamics observed in experiments can be predicted with the precise knowledge of the way individuals use and combine private and social information.

## Experimental Design

We study the individual and collective performance of groups of 5 individuals in a task where each participant has to find the highest values in a 15 × 15 table of 225 cells, each containing a hidden value (see Fig. 1*A*). Supplementary Fig. 1*A* presents an example of a table used in our experiments, where the cell values are explicitly shown. Numbers with values ranging from 0 to 99 were randomly distributed in the cells of the table, and Supplementary Fig. 1*B* shows the distribution of these cell values. To carry out these experiments, we developed an interactive web application that allows the 5 group members to independently explore the same table (see Fig. 1 *B* and *C*).

**Fig. 1:**
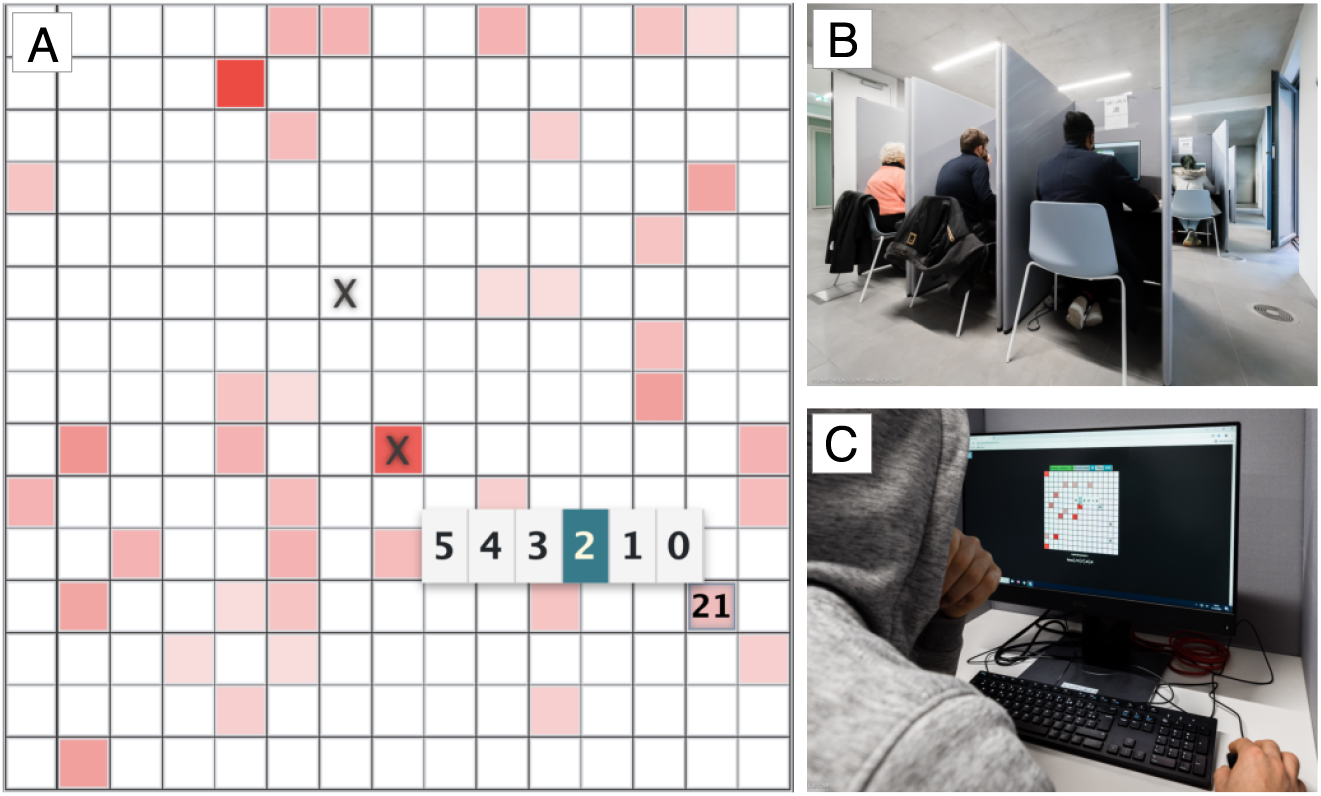
Experimental setup. (*A*) Screenshot of the table at round *t* = 10, as seen by a participant. In this round, the participant has already visited and rated two cells marked with black crosses. He/she has just visited the third cell of value 21 and must rate that cell on a 5-star scale. The score of the participant will then depend on the considered rule: in the non-competitive Rule 1, the score will increase by 0, and by 21 in the competitive Rule 2. (*B*) Pictures of the experimental room and (*C*) of the user interface that participants used during the experiment.

Each experiment includes 20 successive rounds. During each round, each participant has to successively visit and rate 3 distinct cells. Once a participant discovers the hidden value of a cell, he/she must rate that cell on a 5-star scale. The round ends when everyone in the group has visited and rated 3 different cells.

At the start of the next round, the color of each cell in the table is updated according to the percentage of stars that have been used to rate the cell since the beginning of the experiment. The color scale varies between white (0 %) and black (100 %) through a constant gradient of shades of red (see Supplementary Fig. 1*C*). Thus, the cells that have received the highest number of stars since the beginning of the experiment will be clearly visible to all the individuals belonging to the same group. The resulting color map on the table acts like a cumulative long-term collective memory for the group, which is updated at each round. Fig. 1*A* shows an example of a table displaying the participants’ ratings as a color map after 10 rounds during one experiment. Supplementary Fig. 2 illustrates the actions performed by each participant in one group and the color maps associated with the cells in the table resulting from their ratings. Supplementary Movies 1*A* and 2*A* show examples of the dynamics of a typical experiment.

We also investigate the impact of a competitive versus a non-competitive condition on the behavior of participants, and the individual and collective performance. In particular, we focus our analysis on the way individuals visit and rate cells and how they use the traces resulting from their ratings and those left by the other group members to guide their choices. In each experiment, we studied the individual and collective behaviors of two groups performing the same task in parallel and independently. In the non-competitive condition (hereafter called Rule 1), the actions (cell visits and ratings) of the participants do not affect the amount of reward they received at the end of an experiment that always remains constant. On the other hand, in the competitive condition (hereafter called Rule 2), the score of a participant increases at each round by *the value of the cells he/she visits*, but remains unaffected by his/her rating of these cells. Then, the cumulative score of participants over an entire session (12 experiments) determines their monetary reward, which depends on their ranking among the 10 participants and not just among the 5 members of their group (see Materials and Methods for the actual payment method).

This experimental design allowed us to study the impact of an intragroup competition, since each individual in a group competes with the 4 other members of his/her group. However, there is also an implicit intergroup competition, since each individual also competes with the 5 members of the other group for the best rank.

In the next section, we present the results of this experiment mimicking several processes at play in actual 5-star rating systems: *(i)* the exploration by the participants of available options (cells in our experiment), which is greatly influenced by their current ratings; *(ii)* the rating on a 5-star scale of the options selected by the participants, which ultimately affects the future ratings of these different options. The ratings in our experiments, seen by all participants, are digital traces eliciting stigmergic processes allowing the participants to collectively identify the best options. In addition, our basic research study also explores the impact of competition in this exploration/rating context, by submitting the participants to two different incentives (non-competitive/competitive).

Although this competitive aspect is less relevant in real-life situations exploiting 5-rating systems, our experimental setup and our modeling approach allow us, more generally, to study the interplay between exploration strategies, rating strategies, private and shared information, and competition.

## Results

### Collective Dynamics

In this section, we analyze the performance of individuals and groups, as well as the dynamics of collective exploration and ratings in both Rules. To do so, we introduce a set of precise observables, which are described in detail in Materials and Methods: the score of individuals or the mean score of their group; the mean value of the cells weighted by the number of stars or the number of visits at round *t* (*p*(*t*) and *q*(*t*)) or up to round *t* (*P* (*t*) and *Q*(*t*)); the effective number of cells (inverse participation ratio; IPR) over which the stars and visits are distributed at round *t* and up to round *t*; the fidelity *F*, which quantifies whether the distribution of stars or visits in each cell coincides with the actual distribution of the cell values.

Fig. 2 *A* and *B* show respectively the probability distribution functions (PDF) of the score *S* of individuals obtained after the 20 rounds and the score *S*^^^ of groups (i.e., the sum of the scores of the individuals belonging to the same group). In Rule 1, all scores are equal to 0. Thus, in order to compare the individual and collective performance in the two Rules, each individual is assigned a virtual score computed in the same way as in Rule 2. The mean score is higher in Rule 2, showing that this competitive condition provides a stronger incentive to visit high-value cells than in Rule 1: 〈*S/S*_max_〉 = 0.24±0.01 in Rule 1 vs. 〈*S/S*_max_〉 = 0.40 ± 0.01 in Rule 2, where *S*_max_ = 5420 is the maximum theoretical score.

**Fig. 2:**
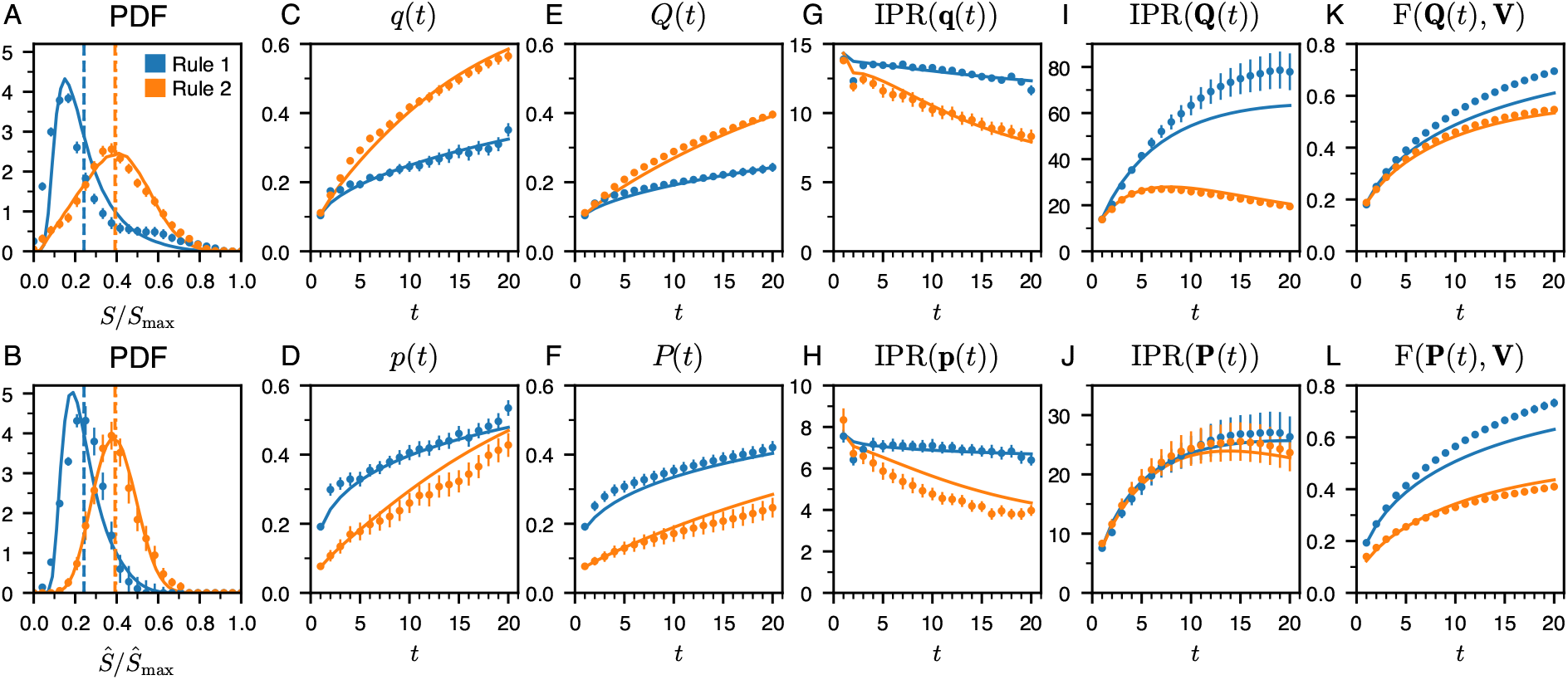
Collective performance and dynamics of collective exploration and ratings. For the non-competitive Rule 1 (blue) and competitive Rule 2 (orange), the symbols correspond to the experimental results, and the solid lines are the predictions of the model. (*A*) Probability distribution function (PDF) of the scores of individuals *S*, and (*B*) of the groups *Ŝ*, respectively normalized by their theoretical maxima *S*_max_ and *Ŝ*_max_ = 5*S*_max_. The dotted vertical lines are the mean score in the experiment, and the dashed vertical lines are the mean scores in the model. (*C*) Average value of the cells visited at round *t*, *q*(*t*) and (*E*) up to round *t*, *Q*(*t*). (*D*) Average value of the cells visited weighted by their ratings at round *t*, *p*(*t*) and (*F*) up to round *t*, *P* (*t*). (*G*) and (*I*) Inverse participation ratio of the visits, IPR(**q**(*t*)) and IPR(**Q**(*t*)), measuring the effective number of cells over which the visits are distributed at round *t* and up to round *t*, respectively. (*H*) and (*J*) Inverse participation ratio of the ratings, IPR(**p**(*t*)) and IPR(**P**(*t*)), measuring the effective number of cells over which the ratings are distributed at round *t* and up to round *t*, respectively. (*K*) Fidelity to the cell value distribution of the distribution of visits, F(**Q**(*t*), **V**), and, (*L*) of ratings, F(**P**(*t*), **V**).

Fig. 2 *C* –*F* show that the average value of the visited cells increases with the number of rounds as the participants discover, visit, and rate cells with higher values. Although *p*(*t*) and *P* (*t*) are higher in Rule 1 than in Rule 2 (Fig. 2 *D* and *F*), the average value of visited cells at round *t*, *q*(*t*), and up to round *t*, *Q*(*t*), are significantly higher in Rule 2 (Fig. 2 *C* and *E*). As we will see later, this apparent paradox is due to the fact that the strategies used by individuals to rate cells in Rule 1 and Rule 2 are very different. In particular, in the competitive Rule 2, some individuals choose to give an average or even a low rating to cells having a high value, presumably to avoid reporting these cells to the other members of their group.

Fig. 2 *G* and *I* show that individuals visit significantly more different cells in Rule 1 than in Rule 2, with IPR(**Q**(*t*)) growing up to the final round *t* = 20 in Rule 1, while it starts decaying after round *t* = 7 in Rule 2. In particular, at the final round *t* = 20 of the experiment, IPR(**Q**(*t* = 20)) is roughly four times larger in Rule 1 than in Rule 2. As we will see in the next section, the lower exploration observed in Rule 2 is mostly due to the fact that the individuals revisit a lot more cells with high values instead of exploring new cells, in order to maximize their score. Moreover, in each round, individuals allocate more stars in Rule 1 compared to Rule 2 (see Fig. 2*H*), but overall, they allocate stars to the same number of cells (see Fig. 2*J*).

Fig. 2 *K* and *L* show that in both conditions, the fidelity increases with the round *t*, suggesting that the correlations between the participants’ visits/ratings and the cell values increase with time. In the final round of the experiment, the fidelity of ratings F(**P**(*t* = 20), **V**) is significantly higher in Rule 1 than in Rule 2. As we mentioned previously, in Rule 1, the participants explore the table a lot more and their ratings better reflect the value of the cells that they have discovered.

### Individual Behaviors

In this section, we characterize the behaviors of individuals and their strategies to visit and rate cells (i.e., the way they use social information in the form of colored traces resulting from their collective past actions), and we quantify the impact of intragroup competition on their behaviors.

#### Choosing the cells to be visited

The probabilities of finding the cells with the highest values are higher in Rule 1 than in Rule 2 (see Fig. 3 *A*–*C* and Supplementary Fig. 3). In Rule 1, individuals find the best cells more often than would be expected if they had searched randomly, illustrating the cooperative effect induced by the use of the digital traces by individuals within groups. In Rule 2, we observe the opposite phenomenon: individuals often revisit the cells that they consider high enough to improve their score without taking the risk of exploring new low-value cells (a sort of hedging), but which also hampers their ability to discover even better cells.

**Fig. 3:**
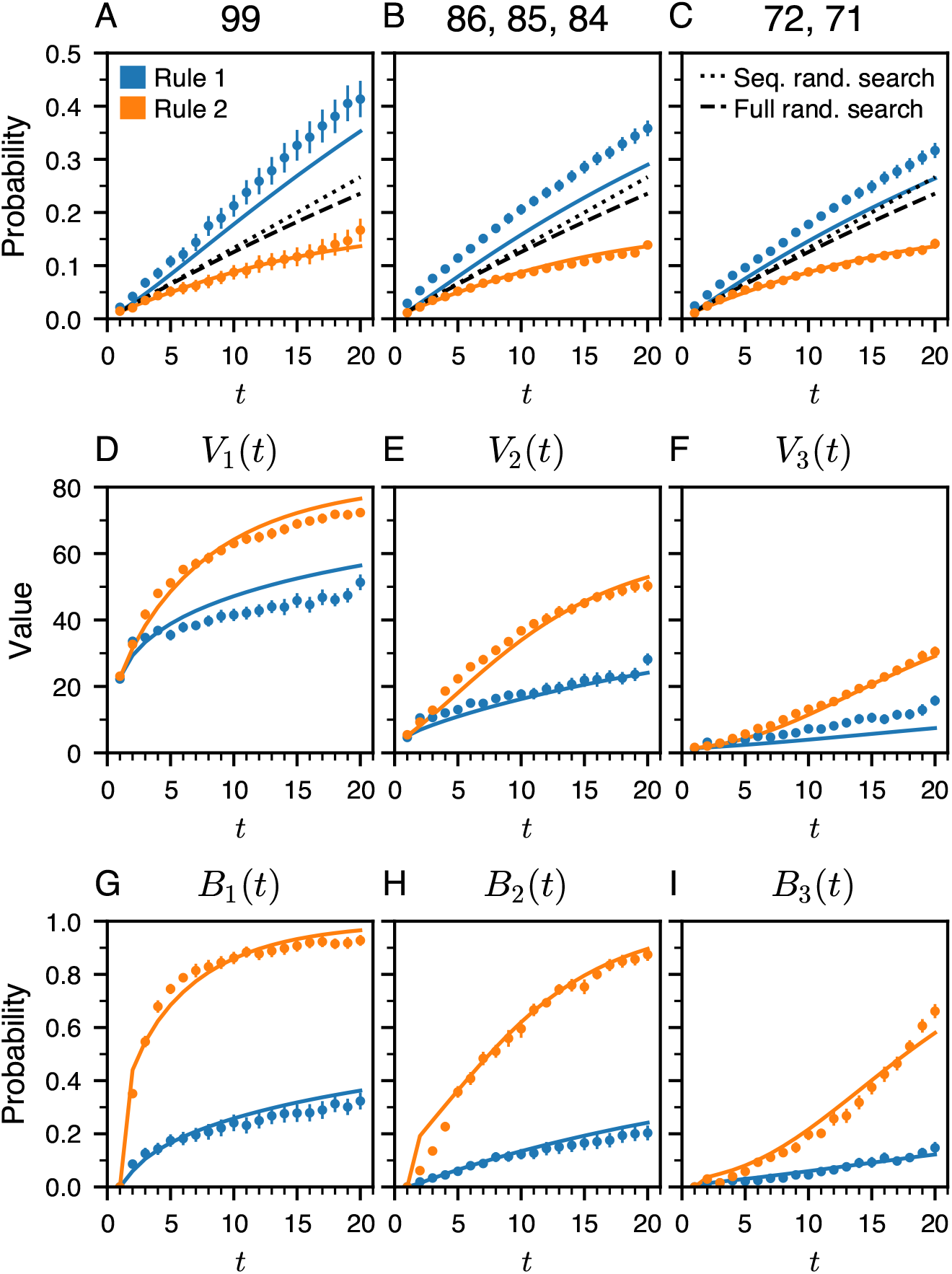
Quantification of individual behaviors for visiting cells. For the noncompetitive Rule 1 (blue) and the competitive Rule 2 (orange), symbols correspond to the experimental results, while solid lines are the predictions of the model. (*A*) Probability to find the best cell, of value 99. (*B*) Probability to find one of the four cells whose values are 86 (× 2), 85, or 84. (*C*) Probability to find one of the four cells whose values are 72 (× 2) or 71 (× 2). The black dashed and dotted lines correspond to the expected probabilities of two different visiting strategies: *i*) cells chosen randomly (full random search, dashed lines), and *ii*) cells chosen randomly among those that have not been already visited (sequential random search, dotted lines). (*D* –*F*) *V*_1_(*t*)*, V*_2_(*t*)*, V*_3_(*t*) are respectively the value of the first-best cell, second-best cell, and third-best cell visited by the participants, as a function of the round *t*. (*G* –*I*) Probability *B*_1_(*t*)*, B*_2_(*t*)*, B*_3_(*t*) to revisit the first-best cell, the second-best cell, and the third-best cell of the previous round, as a function of the round *t >* 1.

We define *V*_1_(*t*), *V*_2_(*t*), and *V*_3_(*t*) as the average of the first-, second-, and third-best values of the cells visited by the participants at round *t*. Fig. 3 *D* –*F* shows that in both conditions, the average values of these 3 cells increase with round *t*. However, their average values are higher in Rule 2. Note that this is not in contradiction with the results shown in Fig. 3 *A*–*C*. As a matter of fact, in Rule 1, individuals have no incentive to revisit cells with high values, so they continue exploring the table even if they have already found those cells. As already mentioned, in Rule 2, individuals have a clear incentive to revisit cells with high values that they can remember, and thus to explore the table less, so that they ultimately discover the cells with the highest values less often.

To confirm this interpretation, we quantify the way individuals revisit cells by defining, for *t >* 1, *B*_1_(*t*), *B*_2_(*t*), and *B*_3_(*t*) as the probability of revisiting at round *t* the cells with the first-, second-, and third-best values of the previous round (*t* − 1). Figure 3 *G* –*I* show that individuals tend to revisit the cells with the best values, and more so as the value of the visited cells increases over time. In the final round of Rule 2, individuals revisit their first-, second-, and third-best cells of the previous round with respective probabilities 93 %, 87 %, and 66 %. In addition, these observables confirm that individuals explore the table more in Rule 1 than in Rule 2: at any round *t* ≥ 5, the values of *B*_1_(*t*), *B*_2_(*t*), and *B*_3_(*t*) in Rule 1 are typically less than one-third of the value in Rule 2.

Altogether, these results illustrate the strong impact of a competitive condition on the way individuals explore the table and select the cells they visit at each round.

#### Rating the visited cells

Supplementary Fig. 4 shows the average fraction of stars *ρ*(*v*) that has been used to rate cells with value *v* at the end of the experiment. *ρ*(*v*) increases with *v*, showing that, on average, individuals give higher ratings to cells having high values and also revisit them more often. The experimental data can be fitted to the following functional form:

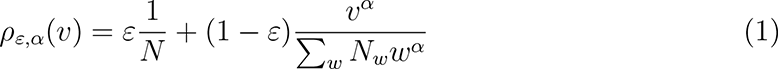

where *ε* ∈ [0, 1] and *α* are two parameters, *N* = 225 is the total number of cells in a table, and *N_v_* is the number of cells with value *v*, such that Σ*_v_ N_v_ρ_ε,α_*(*v*) = 1. Note that the first term *ε/N* quantifies the fraction of stars uniformly deposited in the cells, while the second term involving *α* accounts for the fact that high-value cells should attract more stars.

Supplementary Fig. 5 shows the average number of stars used to rate a cell as a function of its value *v*. In Rule 1, the average number of stars increases almost linearly with *v*. On average, individuals give 1 star to the cells with low values and 4.3 stars to the ones with very high values. In Rule 2 the situation is quite different, individuals give 2.5 stars to low-value cells, and then the average rating decreases to reach a plateau at about 1.5 stars for values higher than *v* = 25. Thus, a cell will receive similar ratings regardless of its value between 35 and 99. This phenomenon suggests that in Rule 2, many participants adopt a non-cooperative/deceptive rating strategy, which effectively makes the information conveyed by the digital trace less discriminating. Overall, these results show that individuals give a much fairer rating to the cells they visit in Rule 1, as the examination of the fidelity has previously revealed.

#### Behavioral profiles of individuals

Supplementary Figs. 6 and 7 show the average number of stars used to rate a cell as a function of its value *v*, for each participant, in Rule 1 and Rule 2, leading to three emerging rating patterns. Some individuals rate cells somewhat proportionally to their value, some rate cells independently of their value, and some others rate cells somewhat inversely proportional to their value.

To quantify and classify these three behavioral profiles, we fit the average rating of each individual with a linear function of the cell value *v*, *u*_0_ + *u*_1_ × 5*v/*99, where *u*_0_ is the intercept and *u*_1_ is the slope of the line (*u*_0_ = 0 and *u*_1_ = 1 would correspond to a linear rating of cells of value 0 to 99, with 0 to 5 stars). Fig. 4 shows the distribution of *u*_0_ and *u*_1_ for all individuals. We identify three classes of behavioral profiles associated with two thresholds at *u*_def−neu_ = −0.5 and *u*_neu−col_ = 0.5 corresponding to the two minima found in the distribution of *u*_1_. These three classes also correspond to those found using Ward’s clustering method on the slope parameter *u*_1_:

**Fig. 4:**
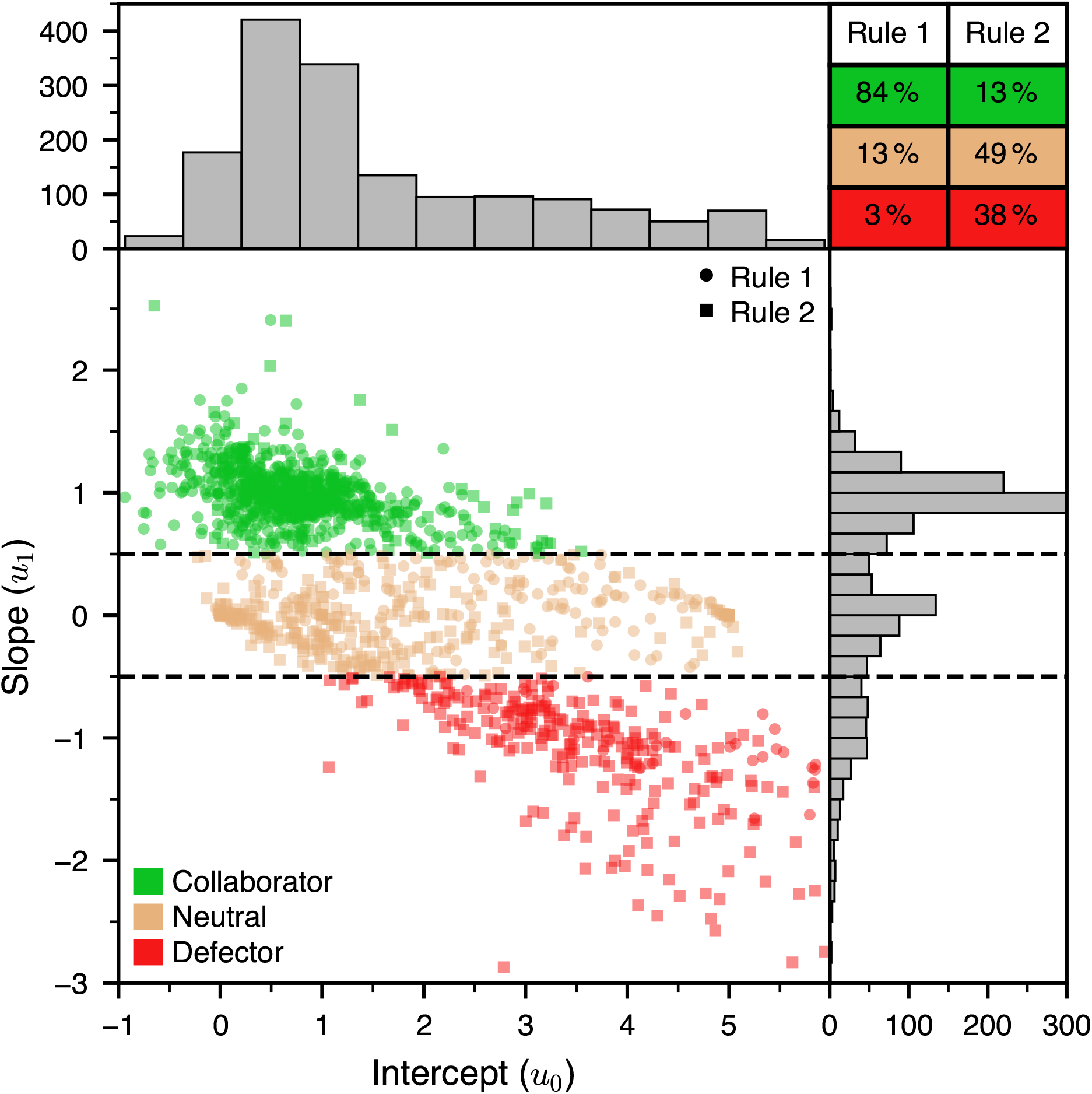
Behavioral profiles of individuals. (Bottom-left) Scatter plot of the values of the two parameters *u*_0_ and *u*_1_ of the linear function, *u*_0_ + *u*_1_×5*v/*99, used to fit each participant’s ratings as a function of the value of the visited cells. In the non-competitive Rule 1, individuals are represented by circles, and in the competitive Rule 2, individuals are represented by squares. The color of the symbols corresponds to the behavioral profile of the individuals: collaborator (green), neutral (brown), and defector (red). The two horizontal lines at *u*_def−neu_ = −0.5 and *u*_neu−col_ = 0.5 are the delimitations between the profiles. (Top-left) Histogram of the values of *u*_0_. (Bottom-right) Histogram of the values of *u*_1_. (Top-right) The table gives the percentage of individuals for each of the behavioral profiles. See also Supplementary Fig. 8A (for Rule 1 only) and *B* (for Rule 2 only).

- Individuals with *u*_1_ ≥ *u*_neu−col_ rate cells in proportion to their values, i.e., they rate cells whose values are the lowest (resp. whose values are the highest) with a small number of stars (resp. a large number of stars; see Fig. 5*A*). Hereafter, we will dub these individuals as *collaborators*, since their rating strategy helps the other members of their group to identify the best cells.
- Individuals with *u*_def−neu_ ≤ *u*_1_ *< u*_neu−col_ rate cells with the same number of stars (on average 3 in Rule 1 and 1.5 in Rule 2) regardless of their values (see Fig. 5*B*). Since the ratings of these individuals do not provide any distinctive information to the other group members, we will dub them as *neutrals*. Note that these neutral individuals do not form a homogeneous group. Indeed, some of them with *u*_0_ close to 0 always give 0 or a very few stars whatever the cell value, hence essentially not participating in the rating and the marking of the cells. Some other neutrals with *u*_0_ close to 5 always give a large number of stars or even 5 stars, thus marking all the cells they visit, while others do not have any consistent logic in the way they rate cells. This explains the wide range of intercepts *u*_0_ ∈ [0, 5] observed for neutrals in Fig. 4. As we will see below, despite not giving distinctive ratings to their visited cells, most neutrals participate in allowing the best cells to emerge by becoming darker on the table, since they often revisit their best-discovered cells.
- Individuals with *u*_1_ *< u*_def−neu_ rate the cells in the opposite way to collaborators, resulting in deceptive ratings. Indeed, they attribute a small number of stars (resp. a large number of stars) to the cells whose values are the highest (resp. whose values are the lowest; see Fig. 5*C*). We will call these individuals *defectors*, since we interpret that the strong traces left on cells with very low values are meant to mislead other group members and prevent them from finding the best cells, especially in Rule 2. In addition, they also decide not to share the position of the best cells they have discovered, by giving them low ratings, and hence not marking them on the table.

Fig. 5 *A*, *D*, and *G* show that collaborators mostly rate cells whose values are less than 20 with 1 star, while the cells whose values are greater than 80 are rated with 5 stars. By contrast, Fig. 5 *B*, *E*, and *H* show that for the neutral individuals, the probability of rating a cell with a given number of stars does not depend on the cell value. Finally, Fig. 5 *C*, *F*, and *I* show that the defectors’ distribution of ratings presents an inverse pattern compared to that of the collaborators. By poorly rating cells with high values (hence hiding them from the other members of their group), while rating cells having low values with a high number of stars (hence misleading others), defectors can then have access to more information than the other group members. Indeed, the defectors benefit from collaborators that only give high ratings to cells having high values. But at the same time, defectors keep private their knowledge about other cells (asymmetrical information [31]) having high values by avoiding marking those cells with a high number of stars while revisiting these cells, thus increasing their scores. Therefore, adopting a defecting behavior can be beneficial in a competitive environment. Indeed, defectors have a higher probability of having the highest score in their group (see Supplementary Fig. 9). However, in the absence of competition, there is no benefit in deception and one should expect fewer defectors. This is what we observe in our experiments, where Fig. 4 (inset table) shows that almost every participant adopts a cooperative behavior in Rule 1, while there is a large fraction of defectors in Rule 2.

**Fig. 5:**
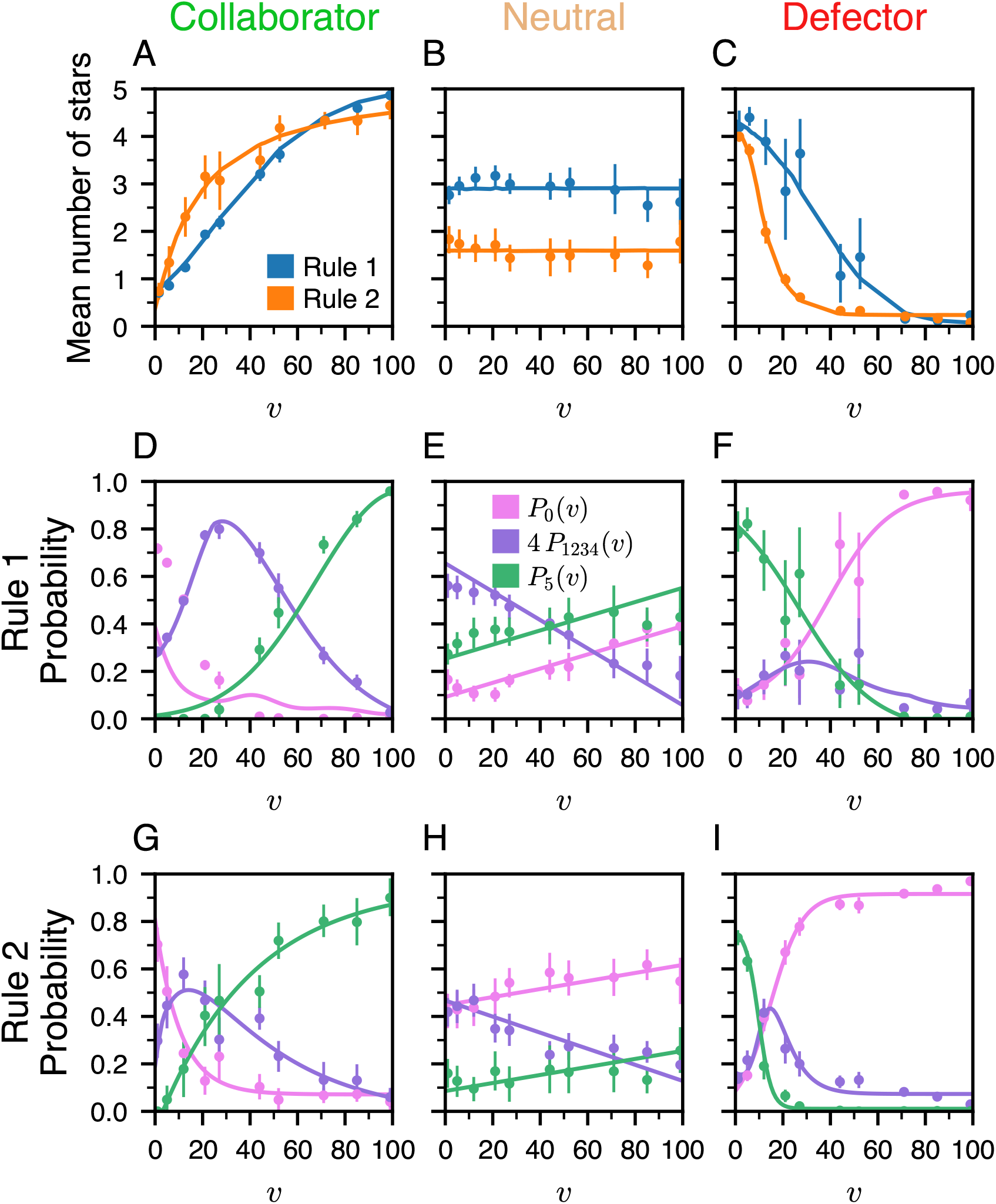
Rating strategies for the three behavioral profiles. (*A*–*C*) Mean number of stars used to rate cells as a function of the cell’s value *v* for (*A*) collaborators, (*B*) neutrals, and (*C*) defectors in the non-competitive Rule 1 (blue) and the competitive Rule 2 (orange). (*D* –*I*) Probability of rating a cell with 0 stars (*P*_0_(*v*); magenta), 1 to 4 stars (*P*_1234_(*v*); violet) and 5 stars (*P*_5_(*v*); green) as a function of its value *v*, for the collaborators, neutrals, and defectors, and for the two rules. The probabilities of rating a cell of value *v* with 1 to 4 stars have been averaged in *P*_1234_(*v*). The dots are the experimental data, and the solid lines are the predictions of the model.

Note that the subjects would participate in 2 experimental runs playing alone (see Supplementary Results) before participating in 10 runs with the 4 other members of their group (in Rule 1 or Rule 2). As expected, when playing alone, the participants behave as collaborators (with themselves), also showing that the players understood well the principle of the game (as confirmed by asking them to fill an anonymous questionnaire at the end of the session).

### Model

We now introduce a stochastic agent-based model to quantitatively identify the strategies for visiting and rating cells, and to understand their respective effects on individual and collective performance. In the model, we simulate groups of 5 agents playing a sequence of 20 consecutive rounds (3 visited and rated cells per round), exactly following the actual experimental procedure. The model, described in detail in Materials and Methods, consists of two steps that characterize the agents’ visit and rating strategies.

The first step accounts for the visit strategy, i.e., which 3 cells an agent decides to visit in each round. This strategy allows for a variety of behaviors observed in the experiment: revisiting the first-, second-, and/or third-best cells already visited in the previous round, depending on their value (private memory; see Fig. 3 *G* –*I*), or exploring a marked or unmarked cell (collective memory; see Supplementary Fig. 4) according to its cumulative fraction of stars (the color of the cell in the actual experiment). The visit strategy is the same for all agents, regardless of their behavioral profile (cooperator, neutral, or defector), as found experimentally, but is allowed to differ for the two conditions, Rule 1 and Rule 2. The second step of the model addresses the rating strategy, i.e., the number of stars an agent uses to rate a visited cell as a function of its value. In the model, the rating strategy of agents depends on their behavioral profile (see Fig. 5 (*D* –*I*)), and is different for the two Rules.

We consider groups of 5 agents (hereafter called MIMIC; see Supplementary Movies 1*B* and 2*B*) that reproduce the behaviors of human collaborators, neutrals, and defectors, and which are drawn according to the fraction of these behavioral profiles observed in the experiments (inset table of Fig. 4). The parameters for the rating strategies of collaborators, neutrals, and defectors have been estimated by fitting the probability to rate a cell with 0 or 5 stars (see Eqs. 5 and 6 in Materials and Methods) to the experimental data (see lines in Fig. 5 (*D* –*I*), and Supplementary Tab. 1). As for the parameters for the visit strategy, they have been estimated by minimizing the error between the experimental and the model results for a set of observables, using a Monte Carlo method (see Supplementary Tab. 2). For all graphs, we ran 1,000,000 simulations, so that the error bars in our simulation results are negligible on the scale of the presented graphs.

Fig. 2 shows that simulations of the model with MIMIC agents quantitatively reproduce the performance of individuals and groups and the observables used to characterize the dynamics of collective exploration and ratings in both Rules, as measured in the experiment. The model also quantitatively reproduces the dynamics of the average value of the first-best, second-best, and third-best cells visited by individuals during the different rounds (Fig. 3 *D* –*F*), along with the probability to revisit each of these 3 best cells at the next turn (Fig. 3 *G* –*I*). In addition, the model reproduces fairly the fraction of collaborators, neutrals, or defectors according to their rank at the end of the experiment and the negative impact of the number of defectors on collective performance (Supplementary Fig. 9). The model also predicts with great accuracy the nontrivial results of Fig. 3 (*A*–*C*), and Supplementary Fig. 3 that were commented above.

These results suggest that the behavioral mechanisms implemented in the model constitute an excellent representation of the processes by which individuals leave and use the traces to guide their choice, and how these processes are modulated in the presence of competition between individuals.

Finally, in the Supplementary Results, we also explore the model predictions for larger group sizes, larger tables, longer durations, and different types of visit and rating strategies. We also consider the optimal parameters of the model that would maximize the score or the ranking of an agent, or the fidelity.

## Discussion

The ability to exploit the traces left in the environment by the action of organisms is one of the simplest and oldest mechanisms used to coordinate collective behaviors in biological systems [32, 33, 34]. In humans, over the past thirty years, the massive development of the Internet, together with applications that extensively use digital traces left voluntarily or not by their users, have reinforced the need to understand how these traces influence individual and collective behaviors [35, 25, 36, 37].

In this work, we have measured and modeled the way groups of individuals leave and use digital traces in an information search task implementing a 5-star rating system similar to the ones used by many online marketplaces and platforms such as Amazon, TripAdvisor, or eBay, in which users can evaluate products, services, or sellers. Although we certainly do not claim that our experimental setup captures all the processes at play in these real-life situations, it shares with them an exploration of the available options (cells in our experiment; products for an online store) greatly influenced by their current ratings, and a rating of the selected options by the participants, allowing the ratings to evolve dynamically.

Our experiment considered two different rules, with Rule 2 implementing a monetary incentive for participants to perform well, resulting in an explicit competition, absent in Rule 1.

Our experimental results show that groups of individuals can use colored traces resulting from their ratings to coordinate their search and collectively find the cells with the highest values in a table of hidden numbers. These traces constitute a form of long-term collective memory of the past actions performed by the group [38, 21]. Combined with the individual short-term memory of the value of the cells already visited, these traces determine the choice of the cells ultimately visited by the participants.

However, our results have also revealed profound disparities in the way individuals use social information resulting from these colored traces to guide them in their tasks, and also in the way they choose to deliver information to other group members through their ratings. We have identified three behavioral profiles (collaborators, defectors, and neutrals) that essentially account for the way in which individuals rate cells. Collaborators cooperate by leaving a trace whose intensity positively correlates with the hidden value of the cells, while the defectors adopt an opposite behavior. Neutral individuals make up a sizable fraction of the group members (13 % in Rule 1 and 49 % in Rule 2) and their ratings are essentially uncorrelated with the actual value of the cells. Yet, the marks that they leave, even if they do not directly inform about the value of the cells, nevertheless induce a cooperative behavior, since neutrals often revisit the high-value cells in a way statistically indistinguishable from the collaborators and defectors.

The information contained in the traces can thus be manipulated by individuals depending on the context, competitive or not, in which the task is performed. Therefore, one may expect that when a situation becomes competitive, individuals should pay less attention to the socially generated traces since the reliability of the information contained in the trace decreases. Previous works in social decision-making have indeed shown that there exists a causal link between mistrust and a decrease in information sharing, and that the fear of being exploited can be a reason for group members to withhold accurate information [39, 40]. This clearly occurs in Rule 2, where 87 % of individuals provide indiscriminate (neutrals) or false (defectors) information, whereas 84 % of individuals (collaborators) provide reliable information in Rule 1.

Despite participants achieving higher scores in the competitive Rule 2 than in Rule 1, by exploring less and revisiting the best cells more, the fidelity of the cumulative trace resulting from their ratings is more faithful to the actual distribution of cell values in Rule 1 than in Rule 2. In other words, there is a better relation (more faithful) between the final rating of a cell and its true value in Rule 1 than in Rule 2, although this relation that we measured remains nonlinear.

We used these experimental observations to build and calibrate a model that quantitatively reproduces the dynamics of collective exploration and ratings, as well as the individual and collective performances observed in both experimental conditions. In par-ticular, this agreement between the model and the experiment is quantified by exploiting a series of subtle observables (PDF of the score, fidelity, IPR, probability of revisiting cells depending on their values. . .). Note that an important added value of our model is to offer (via the analysis of its parameters) a direct and quantitative interpretation of the visit and rating strategies for the three observed behavioral profiles of human participants, and also for different types of optimized agents. The analysis of individual behaviors combined with the simulations of the computational model shows that competition reinforces the weight of private information (i.e., the individual’s memory of the cells already visited) compared to social information (i.e., the collective memory of the group shown on the shared colored table) in the choice of cells that are visited.

The analysis of the model shows that a cooperative effect induced by the trace emerges as soon as there exists a minimal level of marking on cells, and that the fidelity of the ratings increases with cooperation. The model also shows that the trace induces weak cooperation even in groups of defectors, provided they rate cells with a large enough number of stars, simply because they revisit the cells whose values are the highest. In this case, individual memory plays a major role in the collective performance of these defectors. Furthermore, the model predicts that the cooperative effect induced by the traces and the average performance of individuals increases with group size. This property results from the stigmergic interactions between individuals that make it possible to amplify at the group level the information about the location of cells whose values are the highest. Similar properties are observed in many species of ants that use pheromone trail laying to coordinate collective foraging activities and to find the best food sources in their environment [41, 42].

Ultimately, understanding and modeling the processes that govern the influence of social information embedded in digital traces on individual and collective behavior is a crucial step to developing personalized decision-making algorithms as well as artificial collective intelligence systems based on stigmergy [43, 44, 26].

## Materials and Methods

### Ethics statement

The aims and procedures of the experiments were approved by the Ethics Committee of the Toulouse School of Economics (TSE). All participants provided written consent for their participation.

### Experimental procedure

We conducted two series of experiments, the first one in December 2021 to study the competitive condition (Rule 2) and the second one in December 2022 to study the noncompetitive condition (Rule 1). A total of 175 participants were recruited, of which 75 (40 females, 35 males) participated in experiments with Rule 1 and 100 (47 females, 53 males) participated in experiments with Rule 2. Each participant could participate in a maximum of two different sessions. The participants were mostly students at the University of Toulouse, with an average age of 22.

All experiments were carried out at the TSE Experimental Laboratory. After entering the experimental room and before starting the experiment, the participants signed the consent form, were explained the rules, the payment conditions, the anonymity warranty, and were asked to shut down their mobile phones. The participants would then be seated in randomly assigned cubicles (anonymously linked to an ID in our database) that prevented interactions between them (see Fig. 1*B*).

Experiments were conducted using a custom-made interactive web application developed in part in collaboration with the company Andil (www.andil.fr). Participants were presented with the same 15 × 15 table of 225 cells on their respective computer screen, with each cell associated with a hidden value in the range 0 − 99 (examples were provided during the instruction phase). The tables used in the experiments were generated by randomly shuffling the same set of values (see Supplementary Fig. 1*B*). Thus, all tables contained the same set of values, only randomly arranged in the table (see Supplementary Fig. 1*A*).

We conducted a total of 10 sessions with Rule 1 and 15 sessions with Rule 2. At the beginning of each session, each participant performed two consecutive experiments alone (see Supplementary Results for the analysis of these games). The main goal was to ensure that each participant understood the use of the web interface and to measure his/her spontaneous behavior when the only information available was the digital trace resulting from its own activity. Then, the participants were randomly divided into two groups of five and performed 10 successive experiments. During each experiment, the two groups explored different tables that changed during the different experiments.

Each experiment consisted of 20 consecutive rounds, in which each participant had to visit and rate 3 different cells within a recommended time of 20 seconds (beyond which a warning would flash on the screen of late participants). A round would end when all participants in the group had visited and rated 3 cells, and the color of the cells in the table would be updated according to a palette of shades of red that translate the fraction of stars allocated to each cell since the start of the experiment (see Supplementary Fig. 1*C*). participants would then move on to the next round.

In the non-competitive condition (Rule 1), each participant had to find the cells with the highest values in the table, but his/her actions (visiting and rating cells) were not translated into a score. In the competitive condition (Rule 2), the score of each participant would increase at each round by the value of the 3 cells he/she had visited, but it remained independent of the ratings given to these visited cells. Hence, in Rule 2, the participants’ main task was to discover the cells with the highest values, while maximizing their score, and ultimately, their payment at the end of the session. Note that we ultimately introduced a notion of score in Rule 1, to compare the results in the two Rules (see Fig. 2 A and B), although, again, the participants in Rule 1 experiments were never told about any notion of score.

Accordingly, all participants were paid the same 10 Є at the end of a Rule 1 session. In Rule 2, the 10 participants, from the 2 groups of 5, were ultimately ranked and paid according to their cumulated score at the end of the session. The participant ranked first was paid 25 Є, the second was paid 20 Є, the third was paid 15 Є, and the participants ranked from the 4th to the 10th place were paid 10 Є each.

### Observables used to quantify the collective behavior

We define *p_c_*(*t*) as the fraction of stars received by a cell *c* at round *t*. The set of *p_c_*(*t*) for all cells *c* forms a vector **p**(*t*) of size 225 (vectors are shown in boldface). Another vector of interest is the vector **P**(*t*) of the cumulated fraction of stars *P_c_*(*t*) that have been attributed to each cell from the beginning up to round *t* included. Similarly, **q**(*t*) and **Q**(*t*) are vectors whose coordinates *q_c_*(*t*) and *Q_c_*(*t*) represent the fraction of visits received by each cell at round *t* and up to round *t*, respectively.

From the definition of *p_c_*(*t*) and *P_c_*(*t*), we can define the average value of cells visited by the participants weighted by their ratings (number of stars) at round *t*, *p*(*t*) = Σ*_c_ p_c_*(*t*)*V_c_/v*_max_1__, where *v*_max_1__ = 99 is the highest value of a cell. In general, we have *p*(*t*) ≤ 1, and *p*(*t*) = 1 would correspond to all members of a group only giving a non-zero number of stars to the cell of value 99 at round *t*. Similarly, we define the cumulated quantity, *P* (*t*) = Σ*_c_ P_c_*(*t*)*V_c_/v*_max_1__, the average value of cells visited by the participants weighted by their ratings (number of stars) up to round *t*. Hence, *p*(*t*) and *P* (*t*) quantify the instantaneous and cumulated distribution of stars in relation to the value of the visited cells. In particular, a high value of *P* (*t*) (in particular at the final round *t* = 20) indicates that the participants concentrate the allocation of stars on high-value cells. Conversely, a low value of *P* (*t*) suggests a degree of deception, with participants allocating a high fraction of stars to low-value cells, as observed for Rule 2 where many participants are defectors.

In both rules, participants were explicitly asked to discover cells having high values. However, in Rule 2, their score would increase by the value of the cells they visit, thus providing an incentive that affects the way they visit and/or revisit cells during successive rounds. To quantify this (re)visiting behavior, we consider the normalized average value of the cells visited at round *t*, *q*(*t*) = Σ*_c_ q_c_*(*t*)*V_c_* × 3*/*(*v*_max_1__ + *v*_max_2__ + *v*_max_3__), where **V** is the vector of the cell values *V_c_*, and *v*_max_1__, *v*_max_2__, *v*_max_3__ are respectively the first-best, second-best, and third-best values of the cells in the table. This observable is normalized so that *q*(*t*) = 1 corresponds to the best theoretical performance, i.e., when every individual would visit the three best cells of the table at round *t*. Similarly, we introduce *Q*(*t*) that cumulates all visits up to round *t* and which is defined by the same expression replacing *q_c_*(*t*) by *Q_c_*(*t*). Note that, in Rule 2, since the score of the participants is increased by the value of their visited cells, *q*(*t*) and *Q*(*t*) directly quantify the instantaneous and cumulated performance of the group. In Rule 1, the participants had no notion of score, but *q*(*t*) and *Q*(*t*) allow us to characterize the dynamics of their visits, and to compare it with that for Rule 2.

To quantify the exploration behavior of the table by the participants, we introduce the inverse participation ratio (IPR) of the probability vectors **q**(*t*), **Q**(*t*), **p**(*t*), and **P**(*t*). For a given probability distribution **X** = {*X_c_*}, the IPR of **X** is defined as 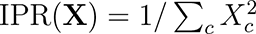. For the 4 vectors considered here, the IPR measures the effective number of cells on which the visits or the ratings are concentrated, at round *t* or up to round *t*. Indeed, if a probability vector **X** is equally distributed over *n* cells among *N*, we have *X_c_* = 1*/n* on these cells, and IPR(**X**) = 1*/*[*n* × (1*/n*)^2^] = *n*, showing that the IPR measures the effective number of cells over which a probability distribution is spread.

We are also interested in the relationship between the hidden values of the cells in the table and the fraction of visits or ratings that these cells have received up to round *t*. This relation is quantified by the fidelity F, which is defined as 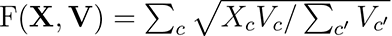, where **X** is **Q**(*t*) or **P**(*t*). The fidelity F takes values in the interval [0, 1] and is equal to 1 if and only if the probability vector **X** is proportional to the vector of cell values **V**, which then corresponds to a perfect fidelity. Indeed, the fidelity can be seen as the scalar product between the vector of coordinates 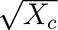 (of unit Euclidean norm, since 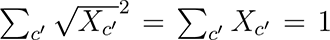) and the normalized vector proportional to 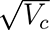. Hence, the fidelity measures how well-aligned these two vectors are and is in fact related to the Hellinger distance between the two distributions. In the context of a real-life 5-star rating system, a high fidelity of the cumulated ratings **P**(*t*) would indicate that the ratings provide a fair representation of the actual value of the different options. Of course, in this context, these intrinsic values of the available options are generally unknown. But our experimental setup provides a simpler context where this relation between the ratings (or the visits) of the different options (the cells, in our experiment) and their intrinsic value (the cell values) can be investigated.

### Model

The agent-based stochastic model includes two components: *(i)* the agents’ strategy for visiting cells; *(ii)* their strategy for rating the visited cells.

#### Visit strategy

In the first round (*t* = 1), the agents have no information, therefore the selection of the 3 cells is fully random. For the other rounds (*t >* 1), the agents adopt the following strategy. For each cell *i* = 1, 2, 3 to visit, they either choose the *i*th-best cell visited in the previous round, of value *V_i_*(*t* − 1), with probability 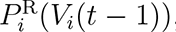, or explore other cells with probability 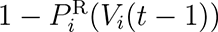, with:

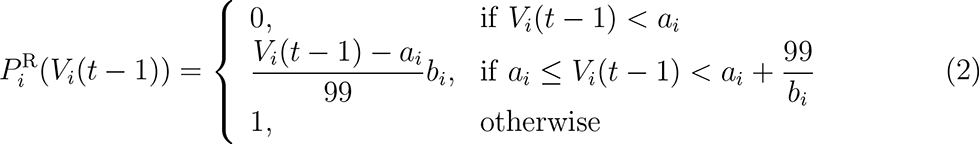

where *a_i_*and *b_i_ >* 0 are parameters. An agent never replays a cell of value *V_i_*(*t* − 1) *< a_i_* and always replays a cell of value *V_i_*(*t* − 1) *> a_i_* + 99*/b_i_* (when this threshold is less than 99, the maximum value of a cell). Between these two thresholds, the probability to revisit the *i*th-best cell linearly interpolates between 0 and 1. The functional form in Eq. 2 is rich enough to be able to capture diverse behaviors, while only using 2 free parameters for each of the 3-best cells, and is in fact consistent with indirect measurements of these probabilities.

When an agent does not visit one of the 3 cells visited in the previous round, it explores other cells in the table. This is done by associating to each cell *c* a probability *P* ^E^(*c, t*) to be selected at round *t*:

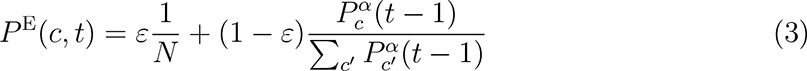

where *P_c_*(*t* − 1) is the fraction of stars deposited in cell *c* up to time *t* − 1, and *ε* ∈]0, 1] and *α >* 0 are parameters. If the selected cell is one of the 3 cells visited in the previous round, another one is selected according to Eq. 3. In Eq. 3, the parameter *ε* controls the amount of exploration of unmarked cells compared to the marked ones: the higher the value of *ε*, the more random the selection (i.e., independent of the cell color). The exponent *α* controls the selection of a cell among the marked ones. A high value for *α* would result in a preferential selection of the highly marked cells, while a small value for *α* would lead to a more homogeneous selection of a cell among the marked ones. The simple functional form in Eq. 3 is inspired by the experimental results of Supplementary Fig. 4, which are well-fitted by the similar functional form in Eq. 1.

The values of the 8 parameters appearing in Eqs. 2 and 3 and characterizing the visit strategy of MIMIC agents in Rule 1 and Rule 2 are reported in Supplementary Tab. 2.

#### Rating strategy

Looking at the probability of rating a cell with *s* stars for each profile (Supplementary Fig. 10), one notes that, except for the collaborators in Rule 1, individuals mostly rate a cell with 0 or 5 stars, and that the other ratings with 1, 2, 3, or 4 stars are less common and have a comparable probability. Therefore, in the model, the probabilities of rating a cell with 1 to 4 stars are set equal and are obtained by imposing the probabilistic normalization condition 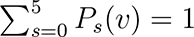, for each value of *v*. In other words, for *s* = 1, 2, 3, 4, we obtain

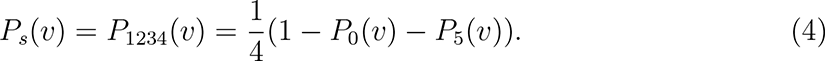

For *s* = 0 and *s* = 5, the probability *P_s_*(*v*) to rate a cell of value *v* with *s* stars is given by

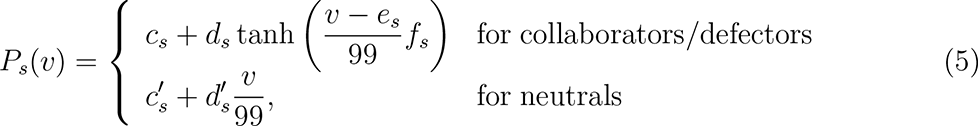

where *c*_*s*_, *d*_*s*_, *e*_*s*_, *f*_*s*_, 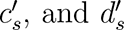 are parameters which must satisfy the property that, for all values of *v*, *P*_0_(*v*) + *P*_5_(*v*) ≤ 1.

However, the *P*_1234_(*v*) approximation is not valid for the collaborators in Rule 1, who use the whole rating scale to rate cells proportionally to their values. Therefore, for these collaborators, we write for *s* = 1, 2, 3, 4, 5,

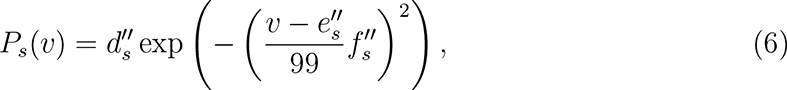

where 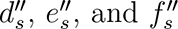 *^′′^* are parameters which must satisfy the property that, for all values of 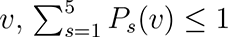. Finally, we set 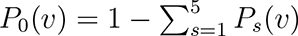.

The functional form of Eqs. 5 and 6 are well adapted to fit the corresponding probabilities observed in the experiment (see lines in Fig. 5 (*D* –*I*) and Supplementary Fig. 10*A*), while allowing to capture very diverse behaviors. Supplementary Tab. 1 presents the values of the parameters appearing in the fitting functional forms of Eqs. 5 and 6.

### Optimization of agents behavior

For the MIMIC agents, the 8 parameters of the visit strategy have been determined by minimizing the error between a set of *n* round-dependent observables, *O*_1_(*t*)*, . . ., O_n_*(*t*), measured in the experiment (by averaging them over every experiment for each of the two considered rules) and the corresponding set of observables, *Ô*_1_(*t*)*, . . ., Ô_n_*(*t*), obtained from extensive simulations of the model (averaging over 1,000,000 numerical experiments for each rule). The error is hence defined by

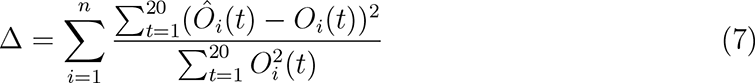

The set of round-dependent observables considered for the computation of this error Δ consists in the following quantities: *q*(*t*), *Q*(*t*), *p*(*t*), *P* (*t*), IPR(**q**(*t*)), IPR(**Q**(*t*)), IPR(**p**(*t*)), IPR(**P**(*t*)), F(**Q**(**t**), **V**), F(**P**(**t**), **V**), *V*_1_(*t*), *V*_2_(*t*), *V*_3_(*t*), *B*_1_(*t*), *B*_2_(*t*), and *B*_3_(*t*). We checked that other sets – in particular, smaller sets – of observables would lead to very comparable results (in particular, in Figs. 2 and 3), fitting some observables slightly better and some others slightly worse, and leading to similar results for the functions characterizing the visit strategy in Eqs. 2 and 3.

To minimize the error in Eq. 7, we have used a Monte Carlo method at zero temperature. At each Monte Carlo step, a small random change is introduced in one of the parameters (also selected randomly). If the error Δ decreases, the new value of the parameter is accepted; otherwise, the old value of the parameter is conserved. The optimization procedure ends when the error stops decreasing. To account for possible multiple local minima of the error, we started the Monte Carlo simulations from several initial values of the parameters. We kept the final parameters, leading to the smallest error. Note that the final parameters obtained in different low-error Monte Carlo runs were found to result in similar functions characterizing the visit strategy in Eqs. 2 and 3.

For the optimization of the parameters of the visit strategy and the parameters of the rating strategy of the optimized agents (see Supplementary Results), we have exploited a similar zero-temperature Monte Carlo method as described above. However, instead of minimizing an error, we have maximized the score or the ranking of the agent, and the fidelity of the traces (see Supplementary Results).

### Computation of the error bars

Error bars for the experimentally measured observables correspond to a level of confidence of 68 % and were determined by exploiting the bootstrap method. Bootstrap is a particular type of Monte Carlo method that evaluates the properties of statistical parameters from an unknown probability distribution by repeated random drawings with replacement from a sample [45]. The bootstrap method starts by creating *M* artificial sets of *N* experiments by drawing with replacement *N* experiments among the *N* original ones. This means that some actual experiments can be drawn more than once in an artificial set, while other experiments may not occur in this set. One can then compute a given observable on every artificial set and obtain its distribution, ultimately leading to confidence intervals. In our case, the independent experiments are the 10 trials played by a group of 5 individuals. Therefore, we have *N* = 20 experiments for Rule 1, and *N* = 15 experiments for Rule 2, and we used *M* = 10, 000 artificial sets to generate bootstrap distributions.

For the numerical simulations of the model, the results correspond to an average of 1,000,000 runs, so that the error bars are negligible on the scale of the presented graphs.

## Supporting information

https://www.dropbox.com/s/01s7svkftdewa1o/SI_Article_Bassanetti_et_al.pdf?dl=0

## Author contributions

C.S. and G.T. designed research; A.B., T.B., S.C., R.E., C.S., and G.T. performed research; M.D. and T.B. designed the web interface, with inputs from all other authors; T.B., C.S., and G.T. analyzed data; T.B. and C.S. designed the model; T.B. performed numerical simulations; T.B., C.S., and G.T. wrote the article.

## Data and materials availability

All data needed to evaluate the conclusions in the paper are present in the paper and/or the Supplementary Materials or available at the following online repository: https://github.com/Thomas-bssnt/Stigmer-article.git.

## Supplementary Materials

Supplementary Results concerning the individuals playing alone and additional model predictions

Supplementary Figures 1 to 20

Supplementary Tables 1 to 3

Supplementary Movies 1 and 2

Supplementary Data 1

