## Supplementary material for "Cooperation and deception through stigmergic interactions in human groups": https://www.dropbox.com/s/01s7svkftdewa1o/SI_Article_Bassanetti_et_al.pdf?dl=0

### Stigmergic cooperation through digital traces in human groups

April 17, 2023

#### This PDF file includes:

Supplementary Results concerning the individuals playing alone and additional model predictions
Supplementary Figures 1 to 20
Supplementary Tables 1 to 3
Supplementary Movies 1 and 2
Supplementary Data 1

#### Other Supplementary Materials for this manuscript include the following:

Supplementary Movies 1 and 2
Supplementary Data 1

### Supplementary Results

#### Behavioral Profiles of Individuals Playing Alone versus in a Group

Before carrying out the experiments in groups, we studied the behavior of the participants playing alone, each individual exploring a different table during two successive rounds and seeing only their own traces (see Supplementary Fig. 11). Supplementary Fig. 12*A* shows that individuals rate the cells similarly to collaborators in groups, except that they rate a low-value cell with 1 star, presumably to remember the cells that they had already opened. Supplementary Fig. 11 *B* and *C* show that in Rule 1, the majority of individuals adopt a collaborative behavior when alone and keep this behavior when they are in a group. On the other hand, in Rule 2, many individuals who adopted a collaborative behavior when playing alone switch to a neutral or defector behavior type when they are in a group.

#### Additional Model Predictions

##### Impact of the number of rounds and group size on individual performance and collective dynamics

Supplementary Fig. 13 shows that after 100 rounds instead of 20 rounds, the normalized score of individuals and groups has increased by 60 % in Rule 2. Beyond round 50, the values of the observables used to quantify the dynamics of collective exploration and ratings begin to saturate. From one round to another, the MIMIC agents revisit almost exclusively the same cells whose values are very high. At the end of the 100 rounds, in Rule 2 the value of their best cell is $V_1(t = 100) \simeq 84$, and the agents revisit their best cell with a probability $B_1(t = 100) \simeq 1$.

Supplementary Fig. 14 shows the impact of group size on the scores of individuals and groups and the dynamics of collective exploration and ratings. We compare the simulation results obtained with groups of 5 MIMIC agents exploring a table of 225 ($15 \times 15$) cells and groups of 20 MIMIC agents exploring a table four times larger, 900 cells ($30 \times 30$). These larger tables were obtained from the combination of four identical tables of 225 cells so that the proportion of each cell value does not change. For instance, in a table of 900 cells, there are four cells with a value of 99, but their proportion (1/225) is the same as in the smaller tables. The dynamics of the inverse participation ratio (IPR) of $\mathbf{p}(t)$, $\mathbf{P}(t)$, $\mathbf{Q}(t)$, and $\mathbf{Q}(t)$ reveal that large groups do not visit four times more cells than small groups, but instead, they concentrate their visits on a few cells with high values. Individuals also have a higher probability of finding the cells with the best values. However, despite these differences, the score remains unchanged. Finally, in Rule 1, the probability that individuals find the best cells at the end of an experiment is much larger in groups of 20 MIMIC agents. Altogether, these results suggest that cooperation induced by stigmergic interactions and the way individuals use the traces resulting from past actions increase with group size.

##### Optimization of agents' performance

We have also exploited our model to find agents that are optimized in different situations. To do this, we used a Monte Carlo method to optimize all the parameters of the model that characterize the visit strategy and the rating strategy.

We first consider a situation in which we maximize the score $S$ (as defined in Rule 2) of five agents (Opt-1 agents) with the same strategy competing in the same group (see Supplementary Figs. 15 and 19*A* and Supplementary Tabs. 1*G* and 2). The inspection of the Opt-1 agents' parameters and Fig. 15 show that they almost rate cells that have very high-values, which they revisit at every round, so that there is almost no exploration. The average score of these optimized agents ($S/S_{\max} = 67\,\%$) is much higher than the score of the participants in Rule 2 ($S/S_{\max} = 40\,\%$).

We then consider a situation in which we maximize the score of one agent exploring a table with four MIMIC agents (see Supplementary Figs. 16 and 19$B$ and Supplementary Tab. 1$H$ and 2). This scenario represents a more realistic situation where an individual seeks to maximize his/her score while competing against four other individuals. In this condition, the behavior of these agents (Opt-2) is markedly different from that of Opt-1 agents, since the presence of MIMIC agents behaving as neutrals and defectors forced them to adapt their visit strategy to cope with indiscriminate or even false information. In terms of rating strategies, the three behavioral profiles result in similar average scores for the agent, but that of MIMIC agents is affected by them, increasing when collaborative and decreasing when behaving as a defector. Nevertheless, the optimization process indicated that the agent with the highest average score is a neutral who assigns 0 stars to every cell. Using these strategies, their average score is $S/S_{\max} = 43\,\%$, which is only slightly better than the one of the average participant.

However, in our experiments, to get the maximum monetary reward, individuals were not required to maximize their score but rather had to optimize their ranking among the two groups of 5 participants. In this condition, the Opt-3 agents behave as defectors (see Supplementary Figs. 17 and 19$C$ and Supplementary Tab. 1$I$ and 2). On average, they obtained a rank of 4.57 among the two groups and a rank of 2.50 within their own group.

Finally, it is interesting to consider the visit and rating strategies maximizing the fidelity of the distribution of ratings to the distribution of cell values in the last round, $\mathrm{F}(\mathbf{P}(t = 20), \mathbf{V})$ (see Supplementary Fig. 18 and Supplementary Tab. 1 and 2). In the long run, the optimal strategy for these agents (Opt-4) is to explore the table randomly and to rate cells proportionally to their value (corresponding to $u_0 = 0$ and $u_1$ in Fig. 4 of the main text). By using this strategy, the agents achieved a fidelity of 0.76 at round 20, and the fidelity would ultimately converge to 1 in the limit of an infinite number of rounds. Obviously, these Opt-4 agents achieve a very mediocre mean score compared to that of the previous optimized agents, and even compared to MIMIC agents reproducing the experimental results. Finally, it is worth noting that there could exist better strategies to maximize the fidelity at round $t = 20$, specifically tailored for the finite 20-round setting used in the actual experiment.

##### Impact of the rating strategy on agents' performance and the fidelity of ratings

To better understand the impact of the rating strategy on individual performance, we studied the collective behaviors of groups of 5 agents having a *linear* rating strategy. These agents rate a cell in proportion to its value, $v$, with $u_0 + u_1 \times 5\,v/99$ stars, where $u_0$ and $u_1$ are respectively the intercept and the slope of the line (see Fig. 4 of the main text). When $u_1 > 0$, the number of stars used to rate a cell increases with its value $v$ (like for a cooperator), while when $u_1 < 0$, the number of stars used to rate a cell decreases with its value $v$ (like for a defector). As $u_0$ increases, agents use a larger number of stars to rate a cell of a given value. Moreover, the combinations of parameters $u_0 \leq 0$ and $u_1 \leq 0$ correspond to a situation in which the agents rate all cells with 0 star, as some actual neutrals do in the experiment. Finally, the visit strategies of these agents are the same as those used by the MIMIC agents in each of the two conditions, Rule 1 and Rule 2.

Supplementary Fig. 20 presents the result of the respective impact of $u_0$ and $u_1$ on ($i$) the average performance of individuals, ($ii$) the average value of cells visited by the participants weighted by their ratings, and ($iii$) the fidelity of ratings with respect to cell values, for each condition Rule 1 and Rule 2.

We first observe that when $u_0 = 0$, as soon as the agents start rating the cells with a non-zero number of stars, the resulting trace allows them to cooperate and significantly increase their performance, even for very low positive values of $u_1$. The results of the simulations also show that the agents get the best scores for negative values of $u_0$, which correspond to situations in which there exists a minimum threshold in the value of a cell that triggers the agents to rate that cell (e.g., when $u_0 = -0.5$ and $u_1 = 0.5$ the threshold is at $v = 20$). Moreover, the higher the value of $u_0$, the worse the performance of the agents. This results from the fact that in that condition, the agents use a very high number of stars with little discrimination in the ratings for different values of $v$. The resulting trace left on cells then provides much less information to the agents, leading to a lower level of cooperation and lower performance. Note however that for high values of $u_0$ (i.e., when $u_0 > 3$) and for weakly negative values of $u_1$ (i.e., when $-1 < u_1 < 0$), there still exists weak cooperation between the agents. At first glance, this is rather counterintuitive, since for these parameters, agents are classified as neutrals or mild defectors. However, this phenomenon can be explained by the fact that, while the traces left by the agents in the initial rounds may not allow for the identification of cells with higher values, over time, cells with higher values will be revisited more often, resulting in a greater accumulation of marks compared to cells with lower values. Nevertheless, for values of $u_1$ that are even more negative, indicating strong defection, the tendency of agents to revisit high-value cells is insufficient to counterbalance the negative impact of assigning high ratings to cells with low values, which ultimately leads to decreased performance.

Finally, the presence of competition between agents (Rule 2) amplifies both the positive and negative effects of the trace compared to the non-competitive situation (Rule 1). Indeed, groups of agents with cooperative behavior ($u_1 > 0$) increase their performance in Rule 2 with respect to the reference situation ($u_0 = u_1 = 0$); conversely, groups of agents with defective behavior ($u_1 < 0$) strongly decrease their performance with respect to the reference situation.

### Supplementary Figures

A

| | | | | | | | | | | | | | | |
|---|---|---|---|---|---|---|---|---|---|---|---|---|---|---|
| 44 | 99 | 0 | 28 | 3 | 0 | 4 | 2 | 1 | 9 | 1 | 0 | 8 | 11 | 1 |
| 2 | 4 | 44 | 2 | 1 | 1 | 3 | 4 | 2 | 7 | 4 | 2 | 3 | 44 | 3 |
| 8 | 2 | 3 | 4 | 2 | 2 | 1 | 1 | 2 | 0 | 1 | 3 | 2 | 45 | 0 |
| 3 | 6 | 2 | 1 | 6 | 2 | 0 | 71 | 3 | 8 | 2 | 3 | 4 | 20 | 20 |
| 0 | 1 | 46 | 85 | 0 | 2 | 43 | 3 | 1 | 1 | 1 | 1 | 6 | 5 | 2 |
| 2 | 2 | 13 | 1 | 0 | 0 | 24 | 86 | 0 | 3 | 2 | 2 | 1 | 3 | 20 |
| 1 | 1 | 20 | 4 | 1 | 2 | 13 | 21 | 22 | 0 | 2 | 2 | 0 | 1 | 6 |
| 3 | 4 | 13 | 1 | 3 | 2 | 1 | 1 | 24 | 6 | 53 | 3 | 3 | 2 | 19 |
| 5 | 3 | 84 | 3 | 2 | 0 | 0 | 3 | 2 | 27 | 3 | 1 | 3 | 14 | 2 |
| 86 | 3 | 2 | 6 | 1 | 45 | 20 | 7 | 24 | 2 | 4 | 27 | 3 | 2 | 3 |
| 1 | 1 | 7 | 0 | 43 | 7 | 1 | 0 | 4 | 2 | 1 | 1 | 53 | 3 | 0 |
| 72 | 2 | 2 | 2 | 6 | 8 | 19 | 3 | 1 | 1 | 72 | 1 | 1 | 1 | 14 |
| 3 | 2 | 4 | 2 | 7 | 0 | 0 | 12 | 0 | 71 | 1 | 3 | 53 | 1 | 9 |
| 2 | 2 | 1 | 8 | 2 | 1 | 0 | 12 | 1 | 2 | 3 | 2 | 1 | 1 | 1 |
| 3 | 1 | 2 | 27 | 51 | 2 | 2 | 3 | 2 | 0 | 2 | 0 | 2 | 3 | 6 |

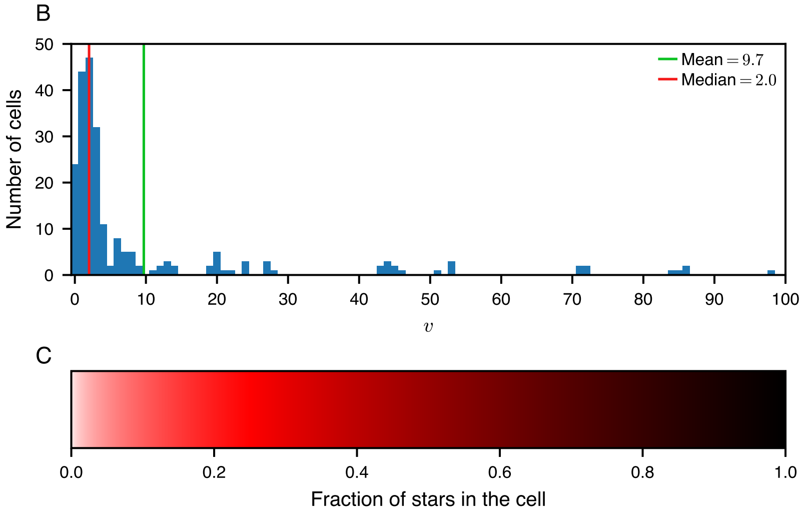


Supplementary Fig. 1: ($A$) Example of a 15 × 15 table used in the experiments and in the simulations of the model (see also Supplementary Movies 1 and 2). ($B$) Distribution of the 225 values $v$ used in the tables. ($C$) Color scale of the visited cells as a function of the fraction of stars used to rate cells since the beginning of an experiment. White color corresponds to cells that have never been visited or to visited cells that have always been rated with 0 stars.

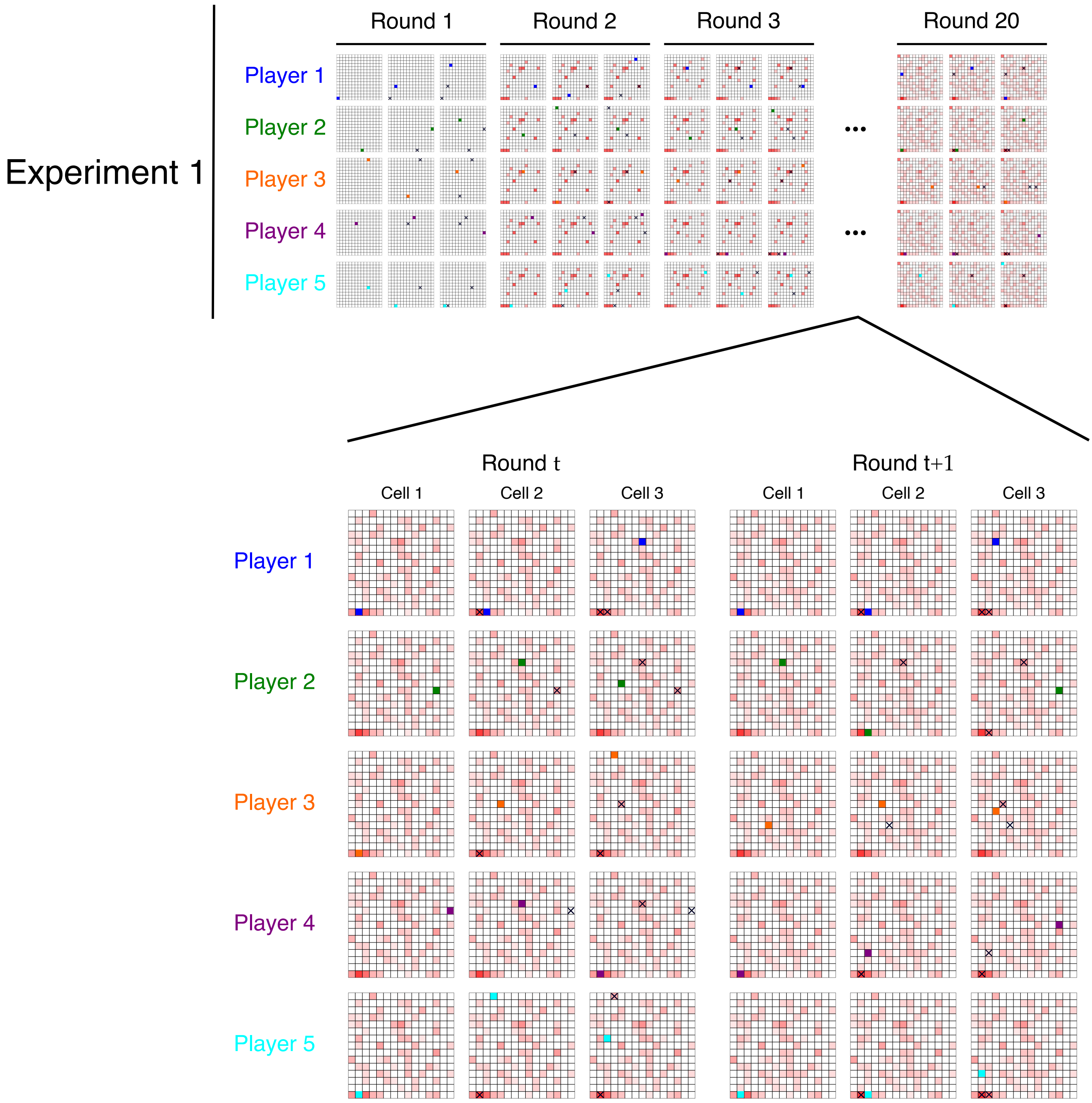


Supplementary Fig. 2: Summary of the experimental protocol. During each round, each participant has to visit and rate successively 3 distinct cells. At the end of each round, the color of each cell in the table is updated according to the percentage of stars that has been used to rate the cell by the five individuals since the first round. The resulting color map on the table acts as a cumulative long-term collective memory for the group.

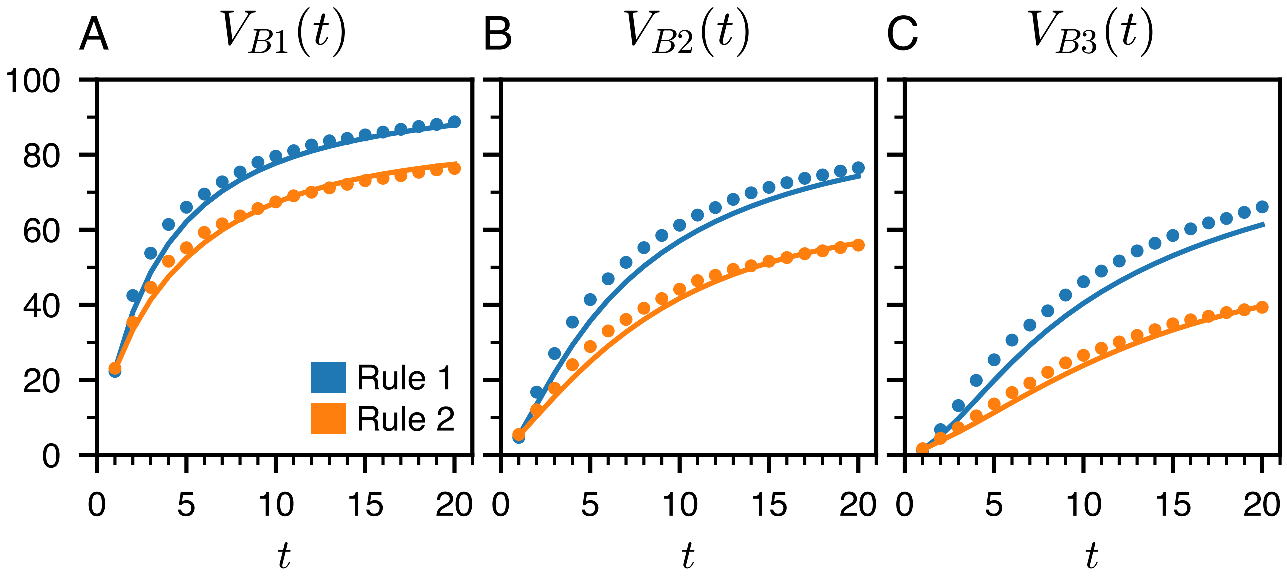


Supplementary Fig. 3: ($A$) First, ($B$) second, and ($C$) third-highest values discovered up to round $t$, as a function of the round $t$, in the non-competitive Rule 1 (blue) and the competitive Rule 2 (orange). The dots are the experimental data, and the solid lines are the predictions of the model.
The highest values discovered are slightly higher in Rule 1 than in Rule 2, showing that the tendency of individuals to revisit cells (and thus to explore less) is higher in Rule 2 than in Rule 1.

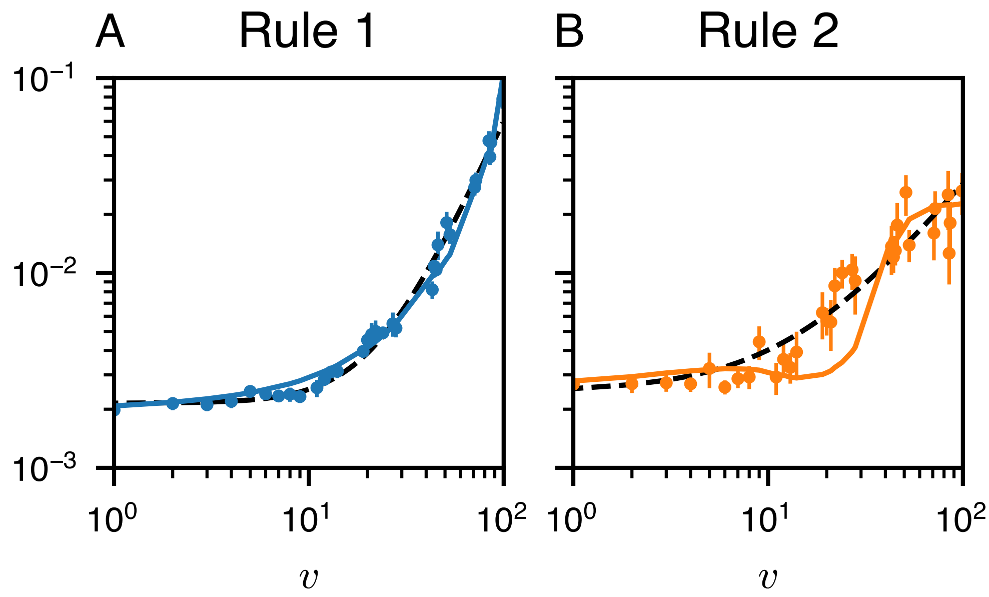


Supplementary Fig. 4: Average fraction of stars $\rho(v)$ used to rate cells of value $v$ at the final round $t = 20$ in Rule 1 ($A$) and Rule 2 ($B$). The black dashed lines correspond to Eq. 1 used to fit the data, with $\varepsilon = 0.48$ and $\alpha = 2.18$ in Rule 1, and $\varepsilon = 0.55$ and $\alpha = 1.22$ in Rule 2.

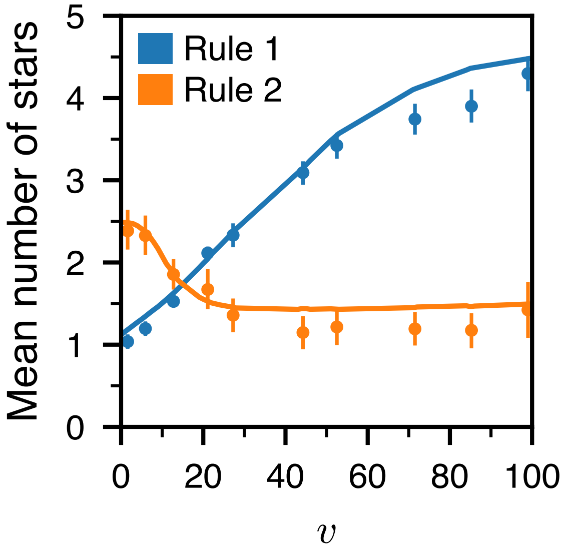


Supplementary Fig. 5: Average number of stars used to rate cells as a function of the cell's value $v$ in the non-competitive Rule 1 (blue) and the competitive Rule 2 (orange). The dots are the experimental data, and the solid lines are the predictions of the model.

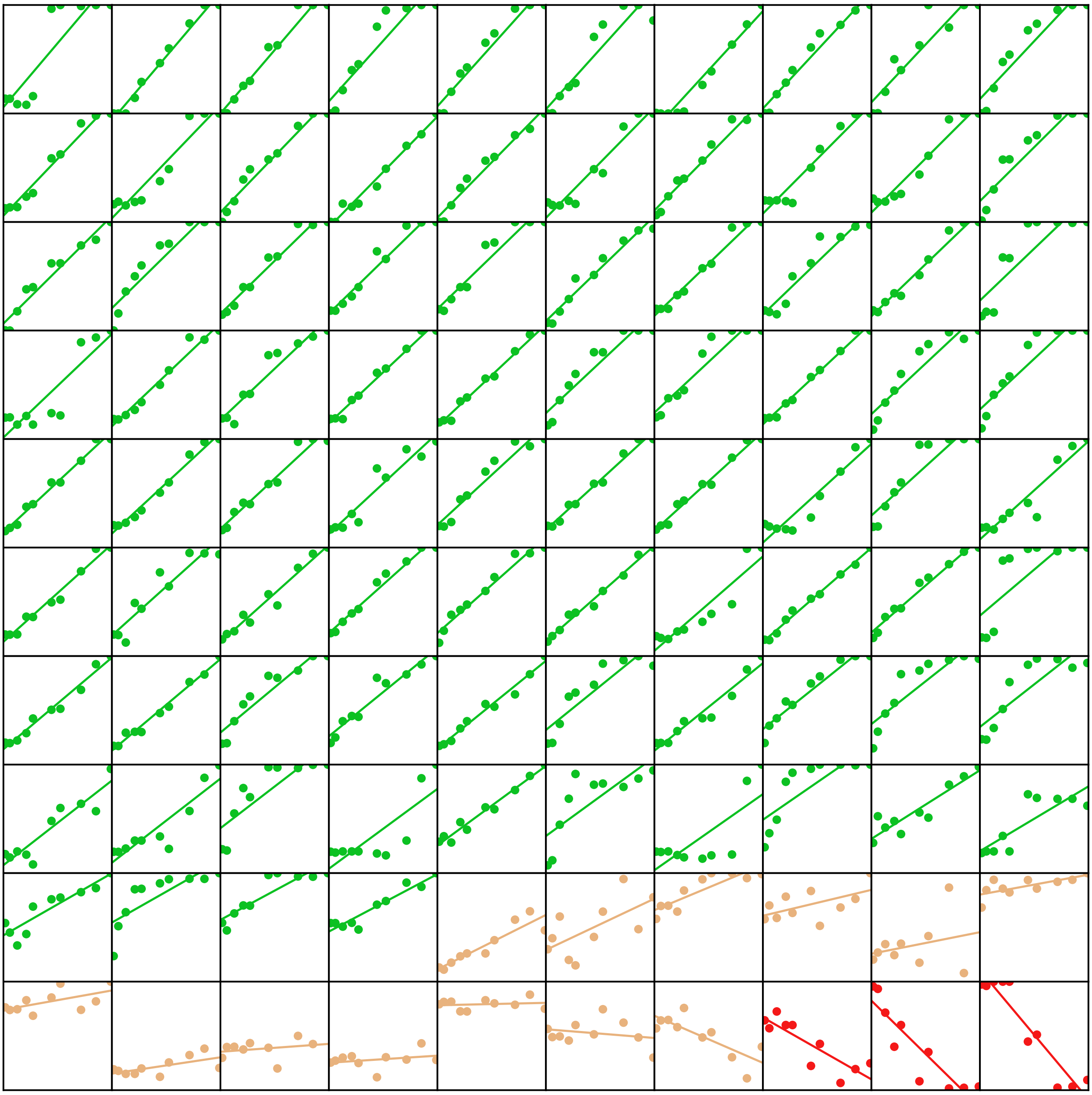

Supplementary Fig. 6: Average number of stars used to rate cells as a function of the cell's value in the non-competitive Rule 1. Each of the rectangles corresponds to the behavior of a single individual aggregated on the 10 experimental runs. The x-axis is the cell's value and goes from 0 to 100 and the y-axis is one-fifth of the number of stars used by the individual to rate a cell of a given value and goes from 0 to 1. The dots are the experimental data, and the line is a linear fit of these data with the function $u_0 + 5\,u_1 v/99$, where $u_0$ is the intercept and $u_1$ is the slope. Individuals are sorted from left to right and from top to bottom according to the value of the slope $u_1$. The color corresponds to the behavioral profile aggregated on the 10 experimental runs: green for collaborators, brown for neutrals, and red for defectors.
Note: Although the individuals' behavior has been defined on each experimental run in Fig. 4 of the main text, we chose to represent the aggregate behavior of each individual averaged over the 10 runs he/she played in a session, in order to limit the number of displayed graphs (100 instead of 1000 if all runs were shown). Therefore, the proportions of each behavioral profile slightly differ from those shown in Fig. 4 (see Supplementary Tab. 3).

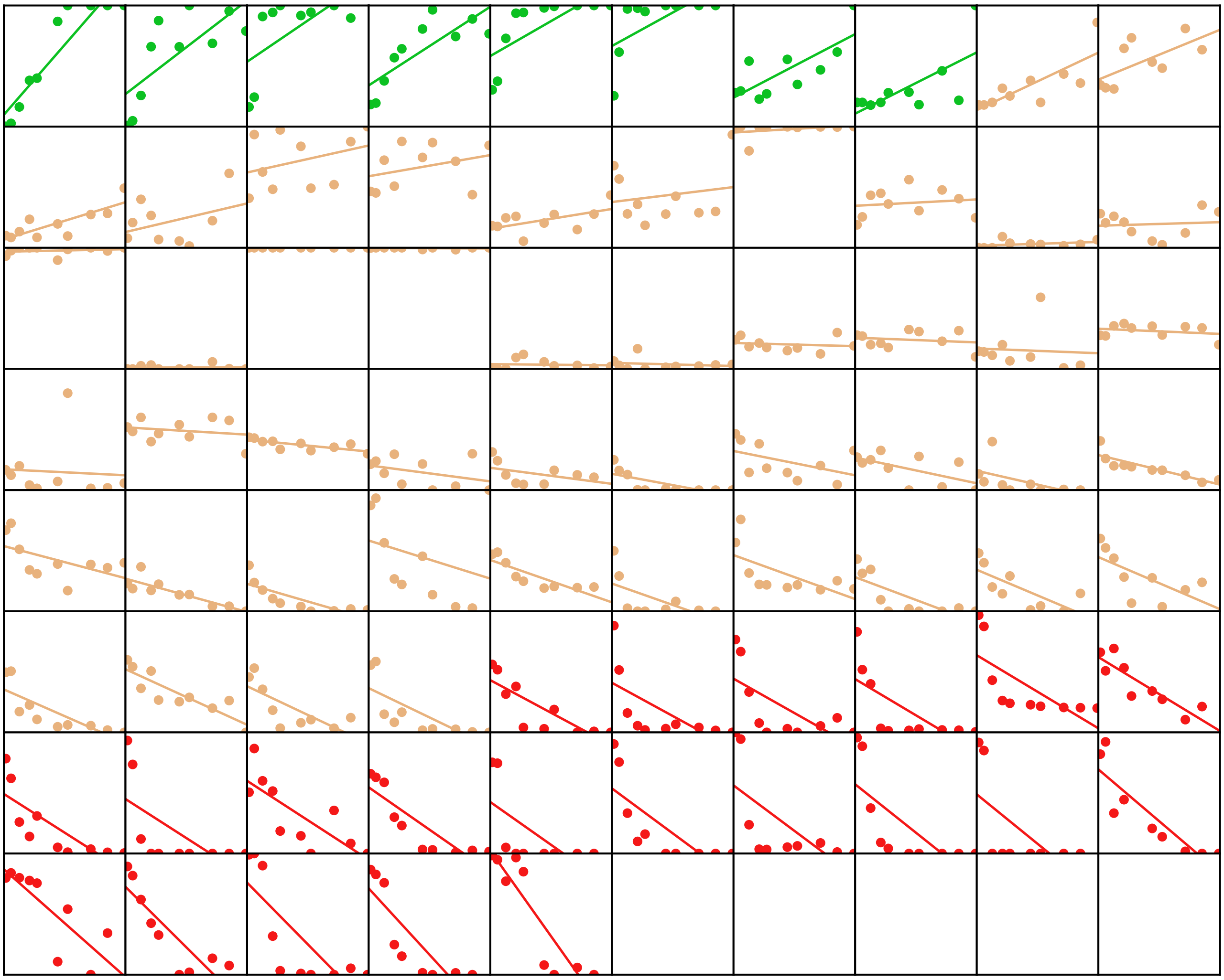

Supplementary Fig. 7: Average number of stars used to rate cells as a function of the cell's value in the competitive Rule 2. Each of the rectangles corresponds to the behavior of a single individual aggregated on the 10 experimental runs. The x-axis is the cell's value and goes from 0 to 100 and the y-axis is one-fifth of the number of stars used by the individual to rate a cell of a given value and goes from 0 to 1. The dots are the experimental data, and the line is a linear fit of these data with the function $u_0 + 5\,u_1 v/99$, where $u_0$ is the intercept and $u_1$ is the slope. Individuals are sorted from left to right and from top to bottom according to the value of the slope $u_1$. The color corresponds to the behavioral profile aggregated on the 10 experimental runs: green for collaborators, brown for neutrals, and red for defectors.
Note: Although the individuals' behavior has been defined on each experimental run in Fig. 4 of the main text, we chose to represent the aggregate behavior of each individual averaged over the 10 runs he/she played in a session, in order to limit the number of displayed graphs (75 instead of 750 if all runs were shown). Therefore, the proportions of each behavioral profile slightly differ from those shown in Fig. 4 (see Supplementary Tab. 3).

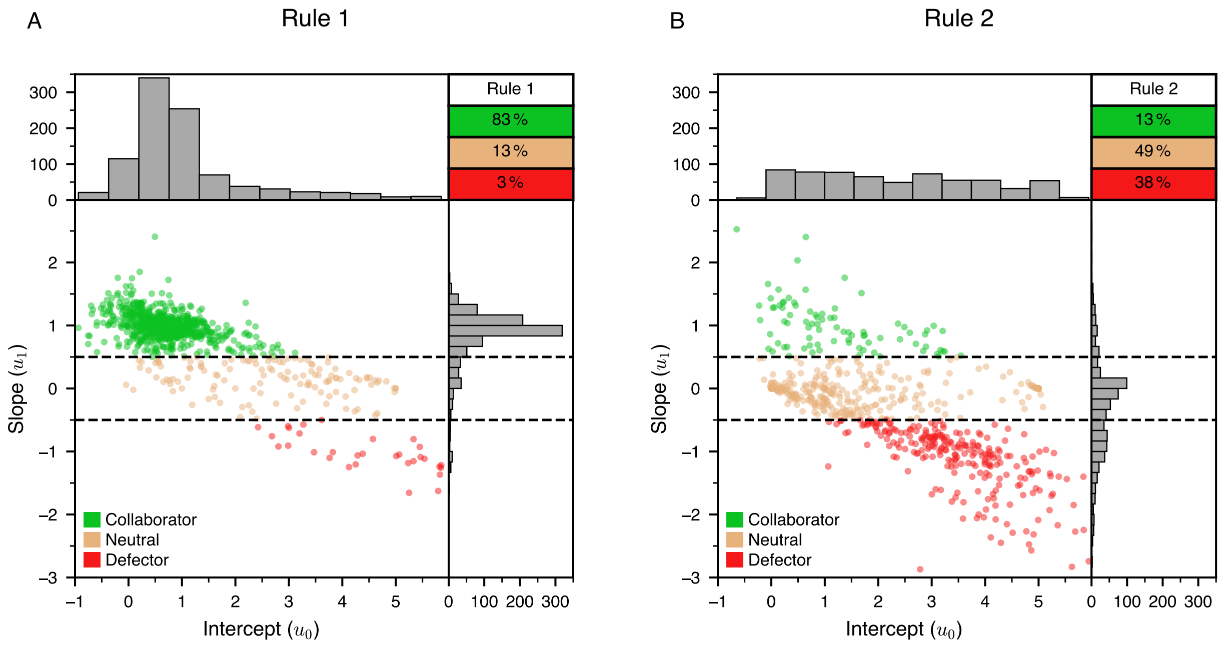


Supplementary Fig. 8: Behavioral profiles of individuals in Rule 1 (*A*) and Rule 2 (*B*). For each subfigure: (Bottom-left) Scatter plot of the values of the two parameters $u_0$ and $u_1$ of the linear function used to fit each subject's ratings as a function of the value of the visited cells. The color of the symbols corresponds to the behavioral profile of the individuals: collaborator (green), neutral (brown), and defector (red). The two horizontal lines at $u_{\mathrm{def-neu}} = -0.5$ and $u_{\mathrm{neu-col}} = 0.5$ are the delimitations between the profiles. (Top-left) Histogram of the values of $u_0$. (Bottom-right) Histogram of the values of $u_1$. (Top-right) The table gives the percentage of individuals for each of the behavioral profiles.

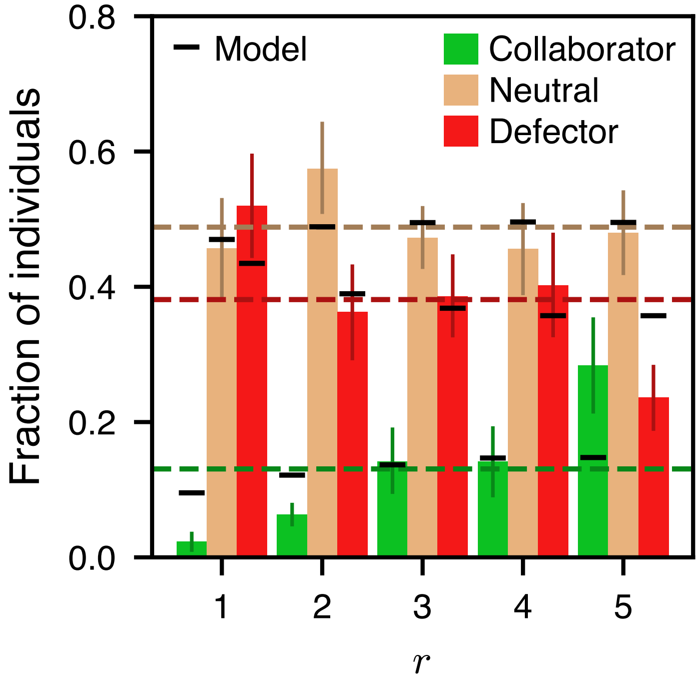


Supplementary Fig. 9: Fraction of individuals with each behavioral profile (collaborator, neutral, and defector) found at ranks $r = 1, 2, \ldots, 5$ (rank determined by their score at the end of the experiment) in Rule 2. The colored bars correspond to experimental data for each behavioral profile: collaborator (green), neutral (brown), and defector (red). The black horizontal lines are the predictions of the model, and the horizontal dashed lines are the proportion of individuals of each behavioral profile in all experiments (null model).
The graph shows that collaborators are less likely to be ranked 1st and more likely to be ranked 5th than expected by the null model, and the opposite is true for defectors. This illustrates the advantage of defectors over collaborators in the competitive Rule 2.

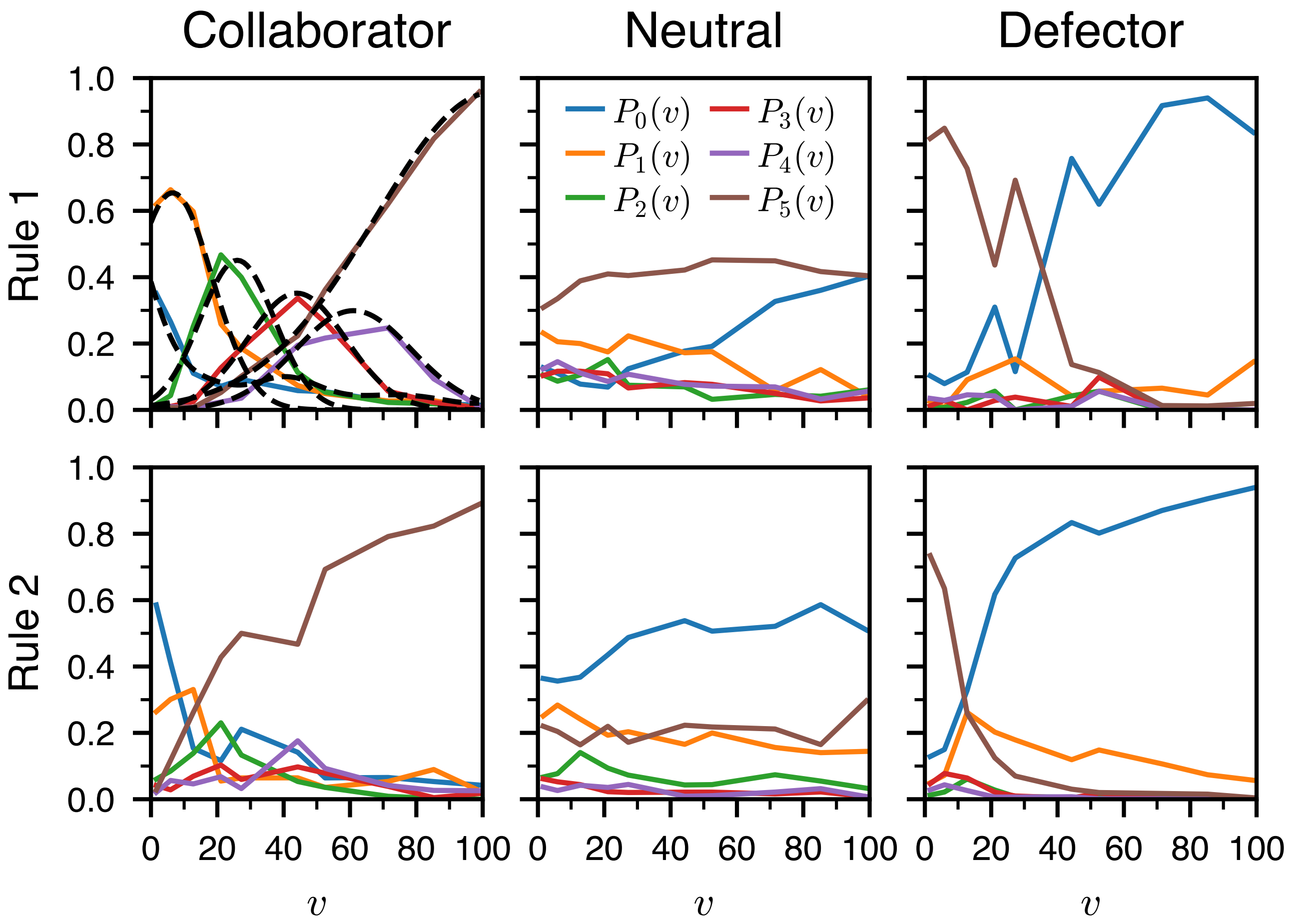


Supplementary Fig. 10: Probability of rating a cell with $s = 0, 1, \ldots, 5$ stars, $P_s(v)$ for the collaborators, neutrals, and defectors, and for the two rules. The solid lines correspond to the experimental data, and the black dashed lines correspond to the fitted Gaussians (Eq. 6) used in the model for collaborators in Rule 1.
Note that $P_0(v)$ and $P_5(v)$ have slightly different values in this figure compared to Fig. 5 *D–I* in the main text. Here, $P_0(v)$ and $P_5(v)$ are the actual experimental values, while in Fig. 5 their values have been slightly adjusted to keep the average number of stars put in a cell of value $v$ unchanged while using the condition $P_1(v) = P_2(v) = P_3(v) = P_4(v) = P_{1234}(v)$.

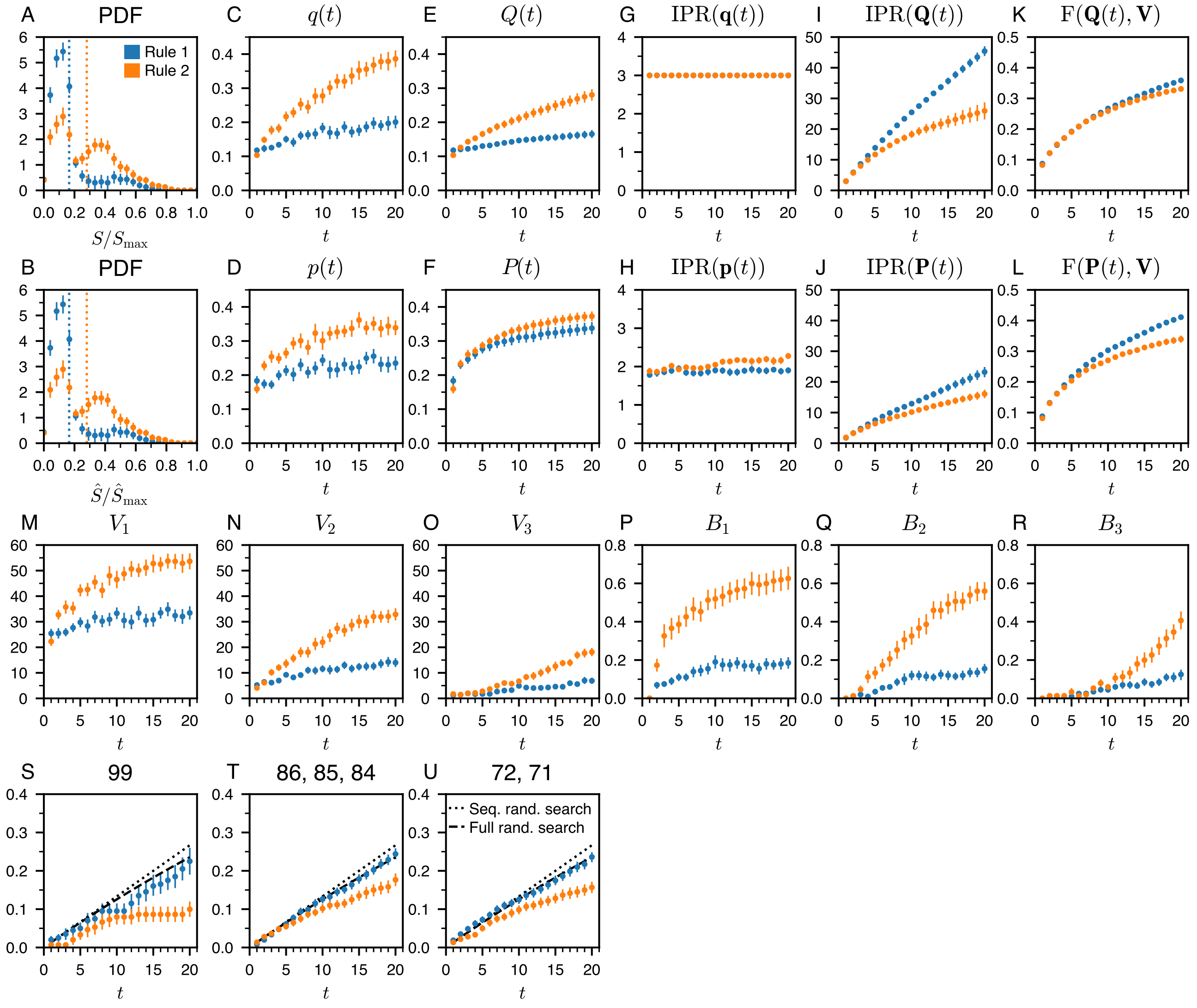


Supplementary Fig. 11: Collective performance and dynamics of collective exploration and ratings for the experiment in which individuals play alone for the non-competitive Rule 1 (blue) and the competitive Rule 2 (orange). ($A$) Probability distribution function (PDF) of the scores of individuals $S$, and ($B$) of the groups $\hat{S}$, respectively normalized by their theoretical maxima $S_{\max}$ and $\hat{S}_{\max} = S_{\max}$. The dotted vertical lines are the mean score in the experiment, and the dashed vertical lines are the mean scores in the model. ($C$) Average value of the cells visited at round $t$, $q(t)$ and ($E$) up to round $t$, $Q(t)$. ($D$) Average value of the cells visited weighted by their ratings at round $t$, $p(t)$ and ($F$) up to round $t$, $P(t)$. ($G$) and ($I$) Inverse participation ratio of the visits, IPR($\mathbf{q}(t)$) and IPR($\mathbf{Q}(t)$). ($H$) and ($J$) Inverse participation ratio of the ratings, IPR($\mathbf{p}(t)$) and IPR($\mathbf{P}(t)$). ($K$) Fidelity to the cell value distribution of the distribution of visits, F($\mathbf{Q}(t), \mathbf{V}$), and, ($L$) of ratings, F($\mathbf{P}(t), \mathbf{V}$). ($M$–$O$) $V_1(t)$, $V_2(t)$, $V_3(t)$ are respectively the value of the first-best cell, second-best cell, and third-best cell visited by the participants, as a function of the round $t$. ($P$–$R$) Probability $B_1(t)$, $B_2(t)$, $B_3(t)$ to revisit the first-best cell, the second-best cell, and the third-best cell of the previous round, as a function of the round $t > 1$. ($S$) Probability to find the best cell, of value 99. ($T$) Probability to find one of the four cells whose values are 86 ($\times$ 2), 85, or 84. ($U$) Probability to find one of the four cells whose values are 72 ($\times$ 2) or 71 ($\times$ 2).

It is worth noting that there are two peaks in the PDF of scores in Rule 2 ($A$). This phenomenon results from the fact that the probability for an individual alone to find a cell with a high-value cell is very low. As a result his/her final score is based solely on exploration.

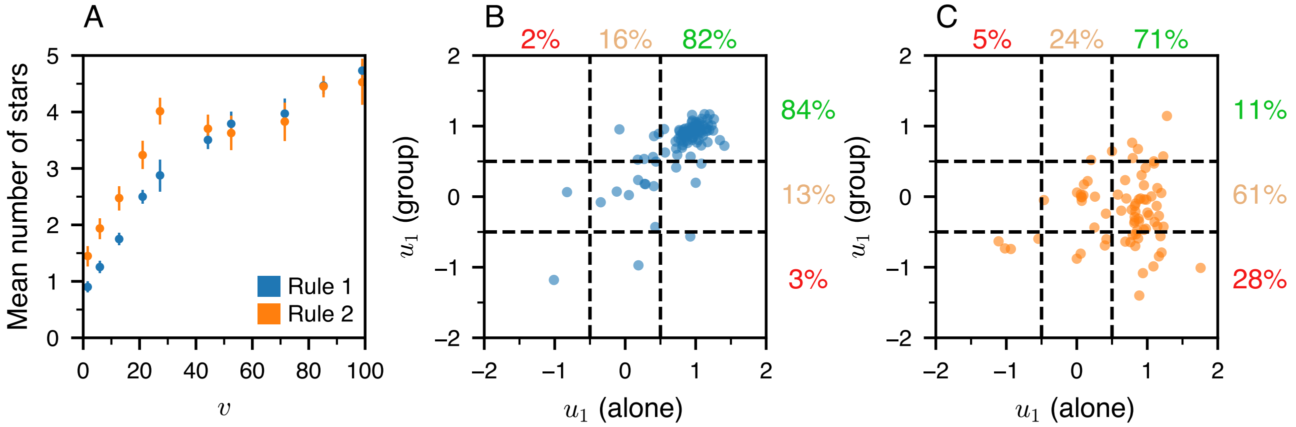


Supplementary Fig. 12: (*A*) Mean number of stars as a function of the cell value $v$ for the experiments in which individuals play alone, for Rule 1 (blue) and Rule 2 (orange). (*B*, *C*) Change in individuals' behaviors between the single-player and five-player experiments. The x-axis represents the average slope $u_1$ of individuals over the two experiments in which they play alone, while the y-axis represents the average slope $u_1$ of individuals over the ten experiments in which they play in groups of five. The two horizontal lines at $u_{\mathrm{def-neu}} = -0.5$ and $u_{\mathrm{neu-col}} = 0.5$ are the delimitations between the profiles. The percentages indicate the fraction of each behavioral profile: collaborators (green), neutrals (brown), and defectors (red).

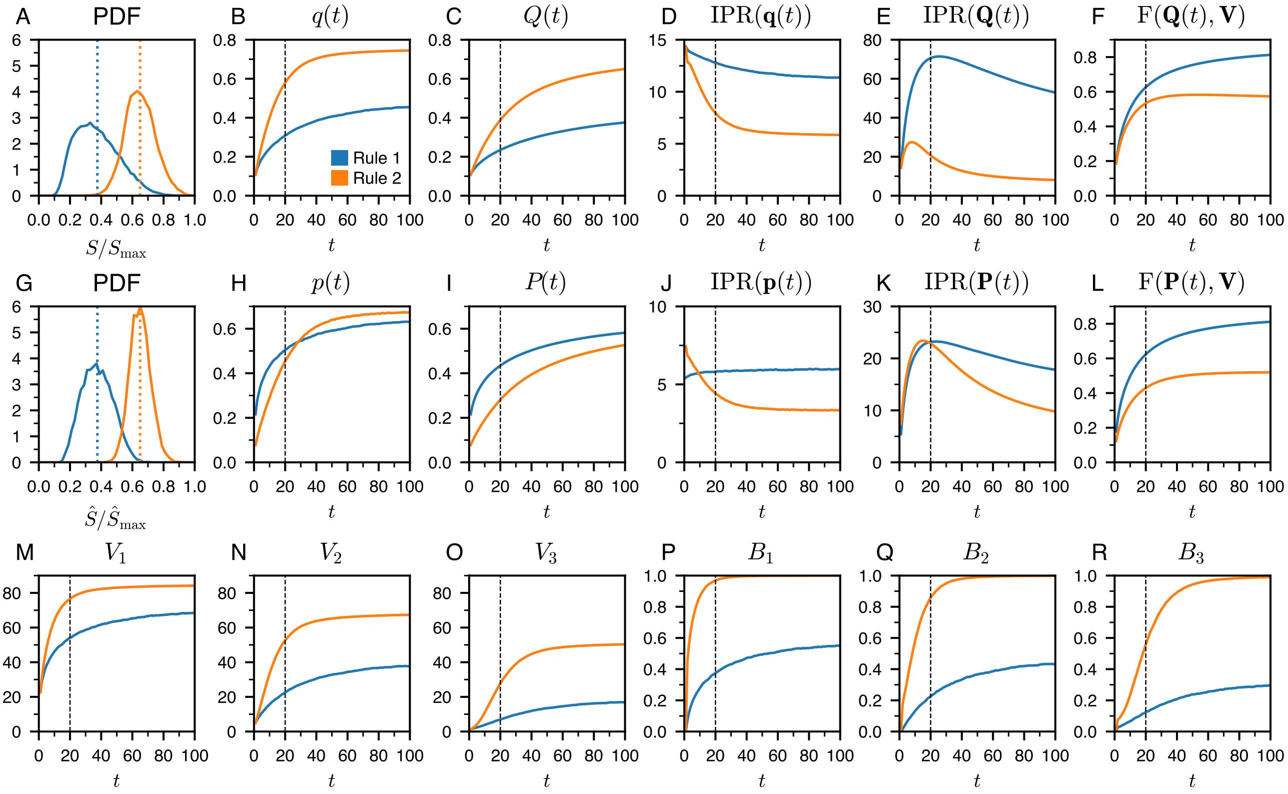

Supplementary Fig. 13: Collective performance and dynamics of collective exploration and ratings in simulations with five MIMIC agents over 100 rounds in Rule 1 (blue), and in Rule 2 (orange). The dotted line at $t = 20$ corresponds to the final round used in the experiments with humans. $(A)$ Probability distribution function (PDF) of the scores of agents $S$, and $(G)$ of the groups $\hat{S}$, respectively normalized by their theoretical maxima $S_{\mathrm{max}}$ and $\hat{S}_{\mathrm{max}} = 5S_{\mathrm{max}}$. The dotted vertical lines are the mean score in the experiment, and the dashed vertical lines are the mean scores in the model. $(B)$ Average value of the cells visited at round $t$, $q(t)$ and $(C)$ up to round $t$, $Q(t)$. $(H)$ Average value of the cells visited weighted by their ratings at round $t$, $p(t)$ and $(I)$ up to round $t$, $P(t)$. $(D)$ and $(E)$ Inverse participation ratio of the visits, $\mathrm{IPR}(\mathbf{q}(t))$ and $\mathrm{IPR}(\mathbf{Q}(t))$. $(J)$ and $(K)$ Inverse participation ratio of the ratings, $\mathrm{IPR}(\mathbf{p}(t))$ and $\mathrm{IPR}(\mathbf{P}(t))$. $(F)$ Fidelity to the cell value distribution of the distribution of visits, $\mathrm{F}(\mathbf{Q}(t), \mathbf{V})$, and, $(L)$ of ratings, $\mathrm{F}(\mathbf{P}(t), \mathbf{V})$. $(M–O)$ $V_1(t)$, $V_2(t)$, $V_3(t)$ are respectively the value of the first-best cell, second-best cell, and third-best cell visited by the participants, as a function of the round $t$. $(P–R)$ Probability $B_1(t)$, $B_2(t)$, $B_3(t)$ to revisit the first-best cell, the second-best cell, and the third-best cell of the previous round, as a function of the round $t > 1$.

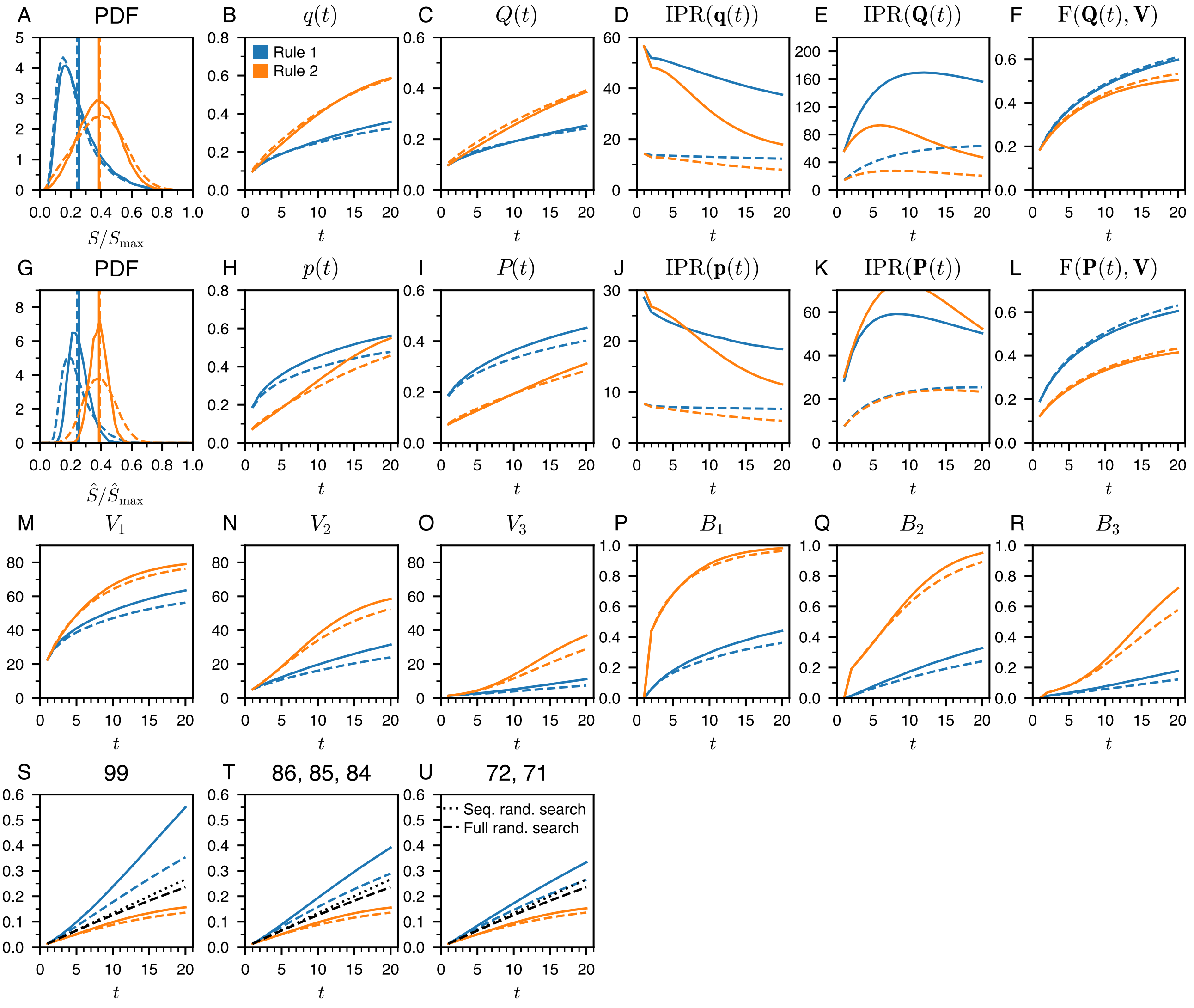


Supplementary Fig. 14: Impact of the group size on the collective performance and the dynamics of collective exploration and ratings in simulations of MIMIC agents for Rule 1 (blue) and Rule 2 (orange). Dashed lines correspond to simulations with five MIMIC agents exploring a table with 225 (15×15) cells, as used in the experiments with humans. Solid lines correspond to simulations with twenty MIMIC agents exploring a table 4 times larger, with 900 (30×30) cells. (*A*) Probability distribution function (PDF) of the scores of agents $S$, and (*G*) of the groups $\hat{S}$, respectively normalized by their theoretical maxima $S_{\max}$ and $\hat{S}_{\max} = 5S_{\max}$ for the dashed line and $\hat{S}_{\max} = 20S_{\max}$ for the solid line. The dotted vertical lines are the mean score in the experiment, and the dashed vertical lines are the mean scores in the model. (*B*) Average value of the cells visited at round $t$, $q(t)$ and (*C*) up to round $t$, $Q(t)$. (*H*) Average value of the cells visited weighted by their ratings at round $t$, $p(t)$ and (*I*) up to round $t$, $P(t)$. (*D*) and (*E*) Inverse participation ratio of the visits, $\mathrm{IPR}(\mathbf{q}(t))$ and $\mathrm{IPR}(\mathbf{Q}(t))$. (*J*) and (*K*) Inverse participation ratio of the ratings, $\mathrm{IPR}(\mathbf{p}(t))$ and $\mathrm{IPR}(\mathbf{P}(t))$. (*F*) Fidelity to the cell value distribution of the distribution of visits, $\mathrm{F}(\mathbf{Q}(t), \mathbf{V})$, and, (*L*) of ratings, $\mathrm{F}(\mathbf{P}(t), \mathbf{V})$. (*M–O*) $V_1(t)$, $V_2(t)$, $V_3(t)$ are respectively the value of the first-best cell, second-best cell, and third-best cell visited by the participants, as a function of the round $t$. (*P–R*) Probability $B_1(t)$, $B_2(t)$, $B_3(t)$ to revisit the first-best cell, the second-best cell, and the third-best cell of the previous round, as a function of the round $t > 1$. (*S*) Probability to find the best cell, of value 99. (*T*) Probability to find one of the four cells whose values are 86 (× 2), 85, or 84. (*U*) Probability to find one of the four cells whose values are 72 (× 2) or 71 (× 2).

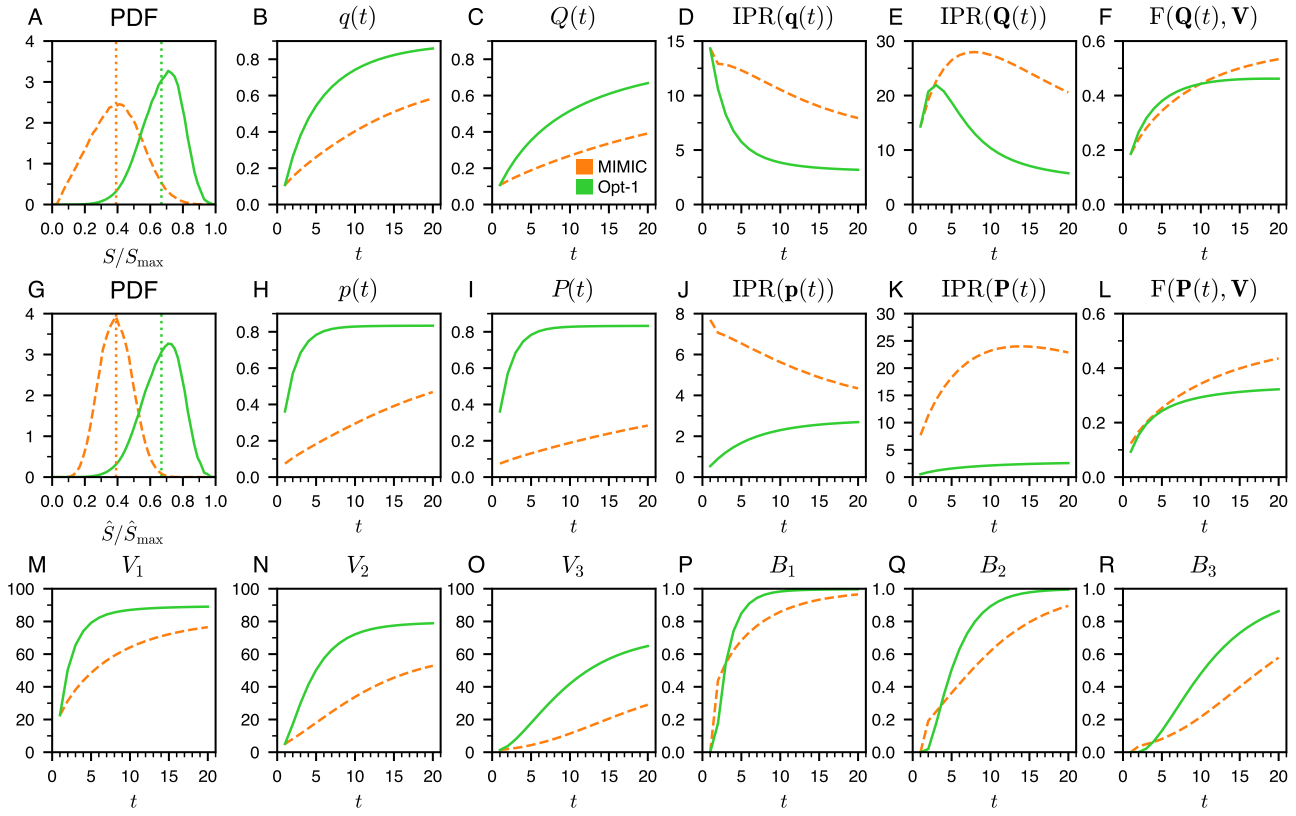


Supplementary Fig. 15: Collective performance and dynamics of collective exploration and ratings in simulations with five Opt-1 agents optimizing the score $S$ (green solid lines) compared to the simulation results with five MIMIC agents (Rule 2, orange dashed lines) which are in good agreement with the experimental results (see Fig. 2 in the main text). ($A$) Probability distribution function (PDF) of the scores of agents $S$, and ($G$) of the groups $\hat{S}$, respectively normalized by their theoretical maxima $S_{\max}$ and $\hat{S}_{\max} = 5S_{\max}$. The dotted vertical lines are the mean score in the experiment, and the dashed vertical lines are the mean scores in the model. ($B$) Average value of the cells visited at round $t$, $q(t)$ and ($C$) up to round $t$, $Q(t)$. ($H$) Average value of the cells visited weighted by their ratings at round $t$, $p(t)$ and ($I$) up to round $t$, $P(t)$. ($D$) and ($E$) Inverse participation ratio of the visits, $\mathrm{IPR}(\mathbf{q}(t))$ and $\mathrm{IPR}(\mathbf{Q}(t))$. ($J$) and ($K$) Inverse participation ratio of the ratings, $\mathrm{IPR}(\mathbf{p}(t))$ and $\mathrm{IPR}(\mathbf{P}(t))$. ($F$) Fidelity to the cell value distribution of the distribution of visits, $\mathrm{F}(\mathbf{Q}(t), \mathbf{V})$, and, ($L$) of ratings, $\mathrm{F}(\mathbf{P}(t), \mathbf{V})$. ($M$–$O$) $V_1(t)$, $V_2(t)$, $V_3(t)$ are respectively the value of the first-best cell, second-best cell, and third-best cell visited by the participants, as a function of the round $t$. ($P$–$R$) Probability $B_1(t)$, $B_2(t)$, $B_3(t)$ to revisit the first-best cell, the second-best cell, and the third-best cell of the previous round, as a function of the round $t > 1$.

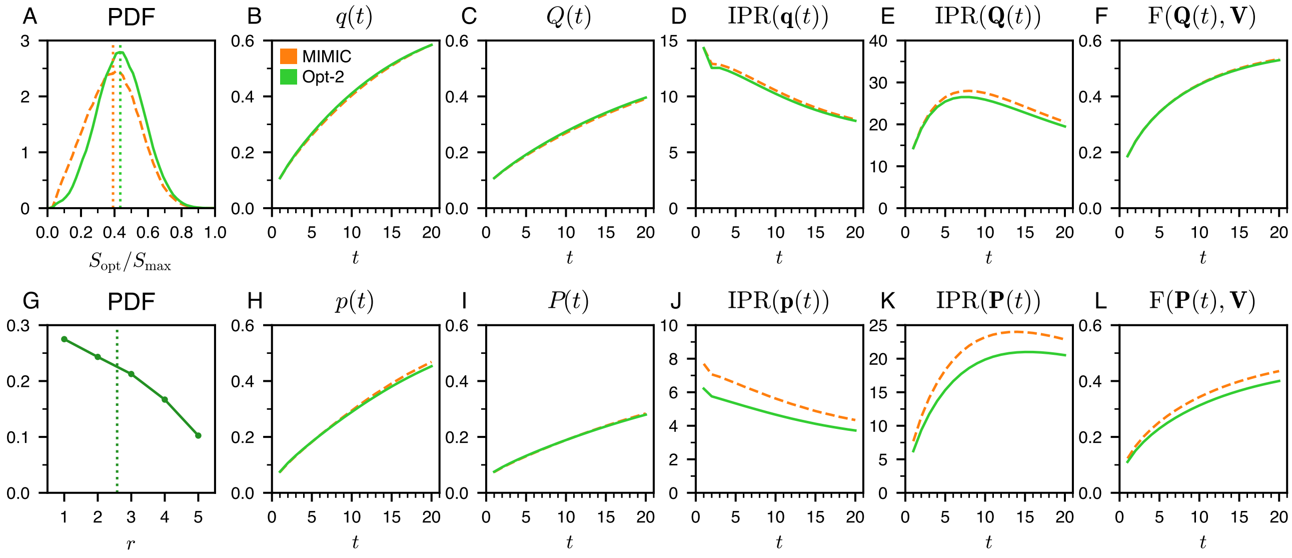


Supplementary Fig. 16: Collective performance and dynamics of collective exploration and ratings in simulations with one Opt-2 agent optimizing its score $S$ playing with four MIMIC agents (green solid lines) compared to the simulations results with five MIMIC agents (Rule 2, orange dashed lines) which are in good agreement with the experimental results (see Fig. 2 in the main text). ($A$) Probability distribution function (PDF) of the scores of agents $S$ normalized by its theoretical maxima $S_{\max}$. The dotted vertical lines are the mean score in the experiment and the model. ($G$) Probability distribution function (PDF) of the rank $r$ of the optimized agent. The dotted vertical lines correspond to the mean rank. ($B$) Average value of the cells visited at round $t$, $q(t)$ and ($C$) up to round $t$, $Q(t)$. ($H$) Average value of the cells visited weighted by their ratings at round $t$, $p(t)$ and ($I$) up to round $t$, $P(t)$. ($D$) and ($E$) Inverse participation ratio of the visits, $\mathrm{IPR}(\mathbf{q}(t))$ and $\mathrm{IPR}(\mathbf{Q}(t))$. ($J$) and ($K$) Inverse participation ratio of the ratings, $\mathrm{IPR}(\mathbf{p}(t))$ and $\mathrm{IPR}(\mathbf{P}(t))$. ($F$) Fidelity to the cell value distribution of the distribution of visits, $\mathrm{F}(\mathbf{Q}(t), \mathbf{V})$, and, ($L$) of ratings, $\mathrm{F}(\mathbf{P}(t), \mathbf{V})$.

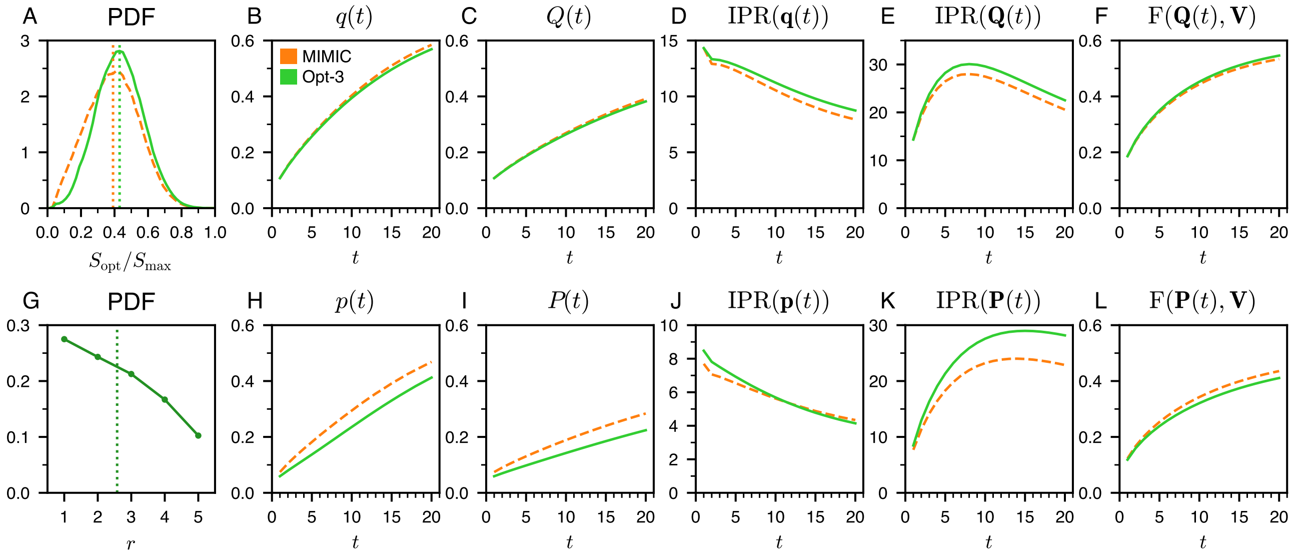


Supplementary Fig. 17: Collective performance and dynamics of collective exploration and ratings in simulations with one Opt-3 agent optimizing its rank $r$ while playing against four MIMIC agents (green solid lines) compared to the simulations results with five MIMIC agents (Rule 2, orange dashed lines) which are in good agreement with the experimental results (see Fig. 2 in the main text). ($A$) Probability distribution function (PDF) of the scores of agents $S$ normalized by its theoretical maxima $S_{\max}$. The dotted vertical lines are the mean score in the experiment and the model. ($G$) Probability distribution function (PDF) of the rank $r$ of the optimized agent. The dotted vertical line corresponds to the mean rank. ($B$) Average value of the cells visited at round $t$, $q(t)$ and ($C$) up to round $t$, $Q(t)$. ($H$) Average value of the cells visited weighted by their ratings at round $t$, $p(t)$ and ($I$) up to round $t$, $P(t)$. ($D$) and ($E$) Inverse participation ratio of the visits, $\mathrm{IPR}(\mathbf{q}(t))$ and $\mathrm{IPR}(\mathbf{Q}(t))$. ($J$) and ($K$) Inverse participation ratio of the ratings, $\mathrm{IPR}(\mathbf{p}(t))$ and $\mathrm{IPR}(\mathbf{P}(t))$. ($F$) Fidelity to the cell value distribution of the distribution of visits, $\mathrm{F}(\mathbf{Q}(t), \mathbf{V})$, and, ($L$) of ratings, $\mathrm{F}(\mathbf{P}(t), \mathbf{V})$.

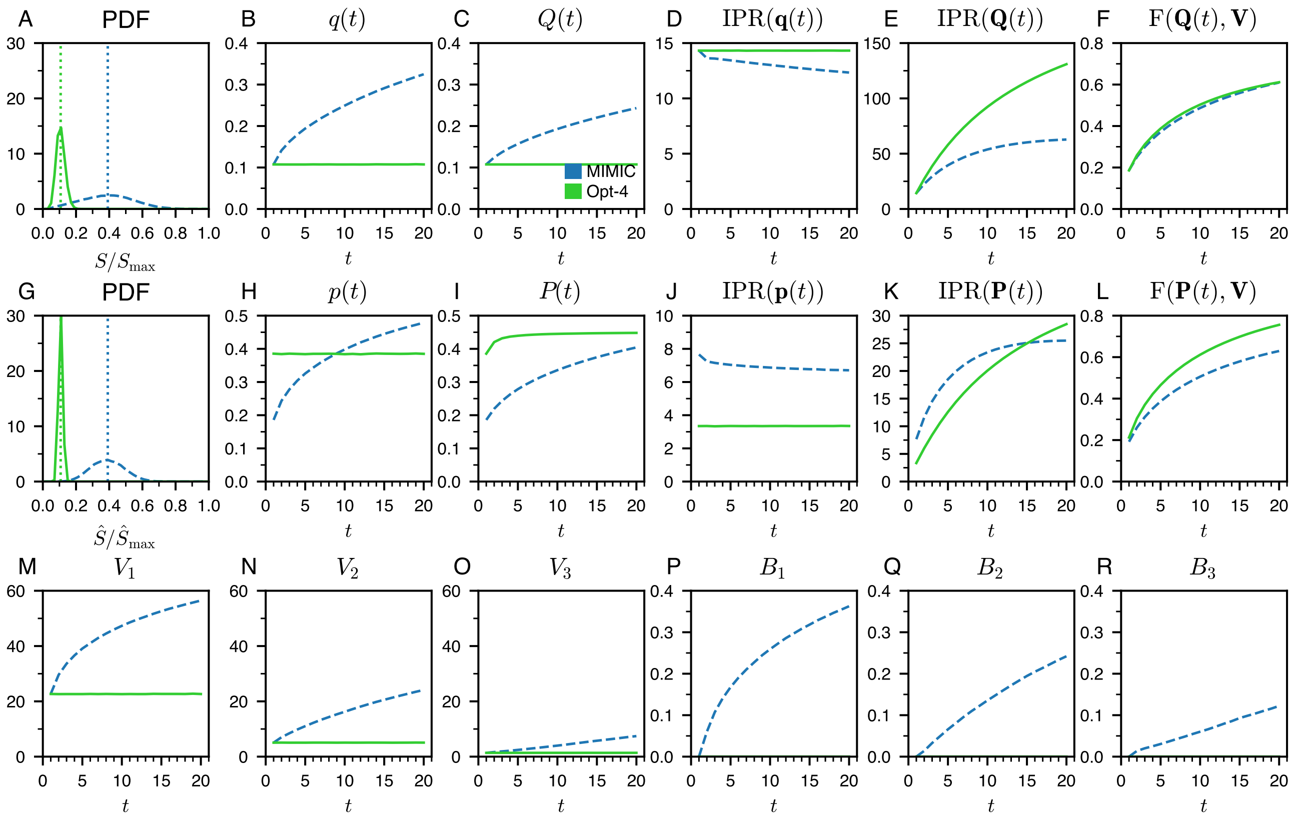


Supplementary Fig. 18: Collective performance and dynamics of collective exploration and ratings in simulations with five Opt-4 agents optimizing the fidelity of ratings with respect to cell values at the end of the experiment $\mathrm{F}(\mathbf{P}(t = 20), \mathbf{V})$ (green solid lines) compared to the simulations results with five MIMIC agents (Rule 1, blue dashed lines) which are in good agreement with the experimental results (see Fig. 2 in the main text). (*A*) Probability distribution function (PDF) of the scores of agents $S$, and (*G*) of the groups $\hat{S}$, respectively normalized by their theoretical maxima $S_{\max}$ and $\hat{S}_{\max} = 5S_{\max}$. The dotted vertical lines are the mean score in the experiment, and the dashed vertical lines are the mean scores in the model. (*B*) Average value of the cells visited at round $t$, $q(t)$ and (*C*) up to round $t$, $Q(t)$. (*H*) Average value of the cells visited weighted by their ratings at round $t$, $p(t)$ and (*I*) up to round $t$, $P(t)$. (*D*) and (*E*) Inverse participation ratio of the visits, $\mathrm{IPR}(\mathbf{q}(t))$ and $\mathrm{IPR}(\mathbf{Q}(t))$. (*J*) and (*K*) Inverse participation ratio of the ratings, $\mathrm{IPR}(\mathbf{p}(t))$ and $\mathrm{IPR}(\mathbf{P}(t))$. (*F*) Fidelity to the cell value distribution of the distribution of visits, $\mathrm{F}(\mathbf{Q}(t), \mathbf{V})$, and, (*L*) of ratings, $\mathrm{F}(\mathbf{P}(t), \mathbf{V})$. (*M–O*) $V_1(t)$, $V_2(t)$, $V_3(t)$ are respectively the value of the first-best cell, second-best cell, and third-best cell visited by the participants, as a function of the round $t$. (*P–R*) Probability $B_1(t)$, $B_2(t)$, $B_3(t)$ to revisit the first-best cell, the second-best cell, and the third-best cell of the previous round, as a function of the round $t > 1$.

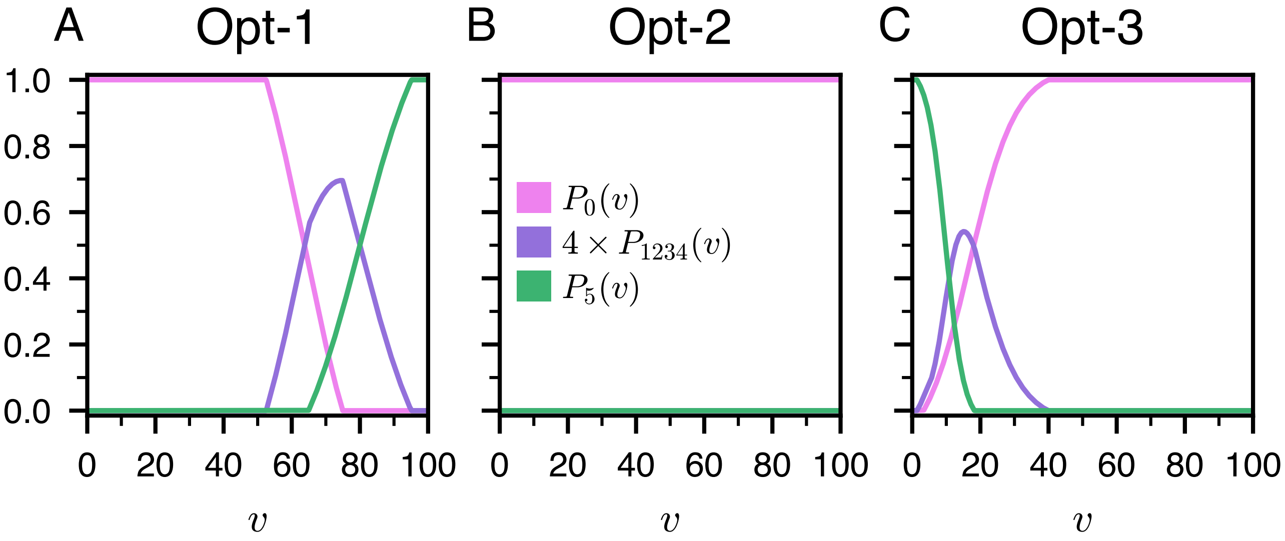


Supplementary Fig. 19: Probability of rating a cell with 0 stars ($P_0(v)$; magenta), 1 to 4 stars ($P_{1234}(v)$; violet) and 5 stars ($P_5(v)$; green) as a function of its value $v$, for the different kind of optimized agents.

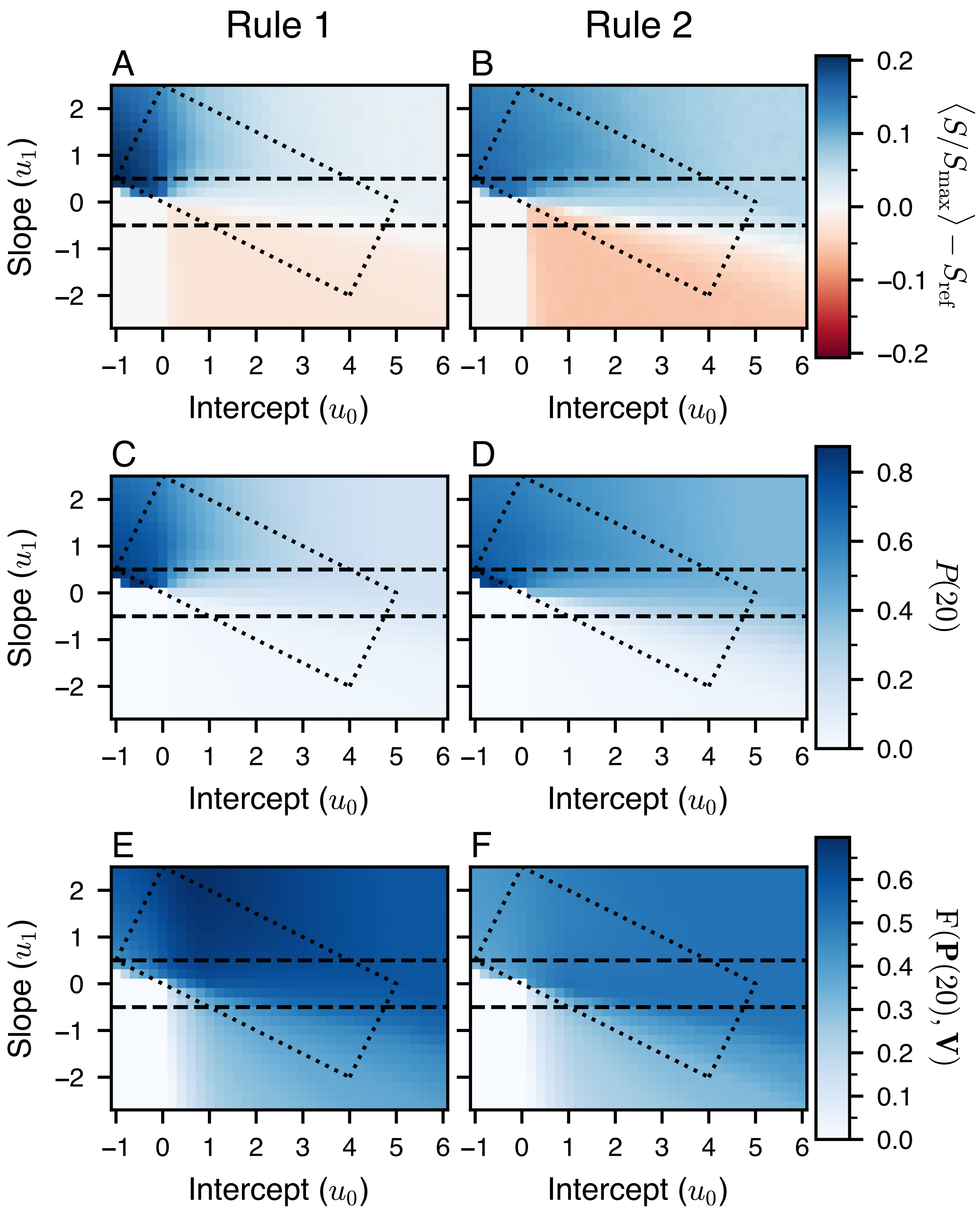


Supplementary Fig. 20: Heatmap for Rule 1 (left column) and Rule 2 (right column) and for different combinations of values of intercept $u_0$ and slope $u_1$ of: $(A)$ and $(B)$ the average value of the score $S/S_{\max} - S_{\mathrm{ref}}$, $(B)$ and $(C)$ the average value of the cells visited weighted by their ratings at the end of the experiment $P(t = 20)$, and $(E)$ and $(F)$ the average value of the fidelity of ratings with respect to cell values at the end of the experiment $\mathrm{F}(\mathbf{P}(t = 20), \mathbf{V})$. Each data point on the heatmap corresponds to the average over 10,000 simulations with five identical agents, defined by their intercept $u_0$ and slope $u_1$. In $(A)$ and $(B)$, $S_{\mathrm{ref}}$ is the normalized score obtained with simulations done with $u_0 = 0$ and $u_1 = 0$. Blue (resp. red) corresponds to positive (resp. negative) values, see color bars. The two horizontal lines at $u_{\text{def-neu}} = -0.5$ and $u_{\text{neu-col}} = 0.5$ are the delimitation between the behavioral profiles, and the rectangle represents the rough location of the agents in the experiments.

### Supplementary Tables

| | $d''_s$ | $e''_s$ | $f''_s$ |
|---|---|---|---|
| $s=1$ | 0.65 | 6.6 | 5.83 |
| $s=2$ | 0.46 | 25.9 | 6.30 |
| $s=3$ | 0.36 | 43.8 | 4.79 |
| $s=4$ | 0.30 | 61.1 | 4.07 |
| $s=5$ | 0.96 | 102.4 | 2.01 |

(A) Collaborator (Rule 1)

| | $c_s$ | $d_s$ | $e_s$ | $f_s$ |
|---|---|---|---|---|
| $s=0$ | 1113.4 | 1113.3 | $-84.5$ | $-4.75$ |
| $s=5$ | $-1051.9$ | 1052.8 | $-304.5$ | 1.24 |

(B) Collaborator (Rule 2)

| | $c'_s$ | $d'_s$ |
|---|---|---|
| $s=0$ | 0.09 | 0.30 |
| $s=5$ | 0.25 | 0.30 |

(C) Neutral (Rule 1)

| | $c'_s$ | $d'_s$ |
|---|---|---|
| $s=0$ | 0.45 | 0.17 |
| $s=5$ | 0.09 | 0.17 |

(D) Neutral (Rule 2)

| | $c_s$ | $d_s$ | $e_s$ | $f_s$ |
|---|---|---|---|---|
| $s=0$ | 0.50 | 0.45 | 39.4 | 3.86 |
| $s=5$ | 0.46 | 0.52 | 26.9 | $-3.11$ |

(E) Defector (Rule 1)

| | $c_s$ | $d_s$ | $e_s$ | $f_s$ |
|---|---|---|---|---|
| $s=0$ | 0.45 | 0.46 | 14.8 | 7.34 |
| $s=5$ | 0.39 | 0.38 | 9.8 | $-18.49$ |

(F) Defector (Rule 2)

| | $c_s$ | $d_s$ | $e_s$ | $f_s$ |
|---|---|---|---|---|
| $s=0$ | 0.5 | 0.95 | 63.7 | $-5.17$ |
| $s=5$ | 0.5 | 0.78 | 80.1 | 5.01 |

(G) Opt-1

| | $c'_s$ | $d'_s$ |
|---|---|---|
| $s=0$ | 1 | 0 |
| $s=5$ | 0 | 0 |

(H) Opt-2

| | $c_s$ | $d_s$ | $e_s$ | $f_s$ |
|---|---|---|---|---|
| $s=0$ | 0.45 | 0.59 | 16.9 | 7.34 |
| $s=5$ | 0.51 | 0.55 | 9.8 | $-18.48$ |

(I) Opt-3

Supplementary Tab. 1: Parameters values used for the rating strategy (see Eqs. 5 and 6 in the main text) for MIMIC agents (collaborator, neutral, and defector) in both rules, and for the optimized agents (Opt-1, Opt-2, Opt-3). These values result from the fitting of the probabilities of rating a cell with $s$ stars described in the main text.

| | | $P^{\mathrm{E}}(c,t)$ | | $B_1(t)$ | | $B_2(t)$ | | $B_3(t)$ | |
|---|---|---|---|---|---|---|---|---|---|
| | | $\varepsilon$ | $\alpha$ | $a_1$ | $b_1$ | $a_2$ | $b_2$ | $a_3$ | $b_3$ |
| Rule 1 | MIMIC | 0.78 | 0.89 | 57.6 | 2.19 | 25.0 | 2.29 | 1.4 | 2.64 |
| Rule 2 | (col, neu, def) | 0.69 | 1.32 | −8.4 | 1.55 | −4.1 | 2.11 | −0.2 | 2.33 |
| | Opt-1 | 1e-5 | 1.38 | 25.0 | 2.00 | 18.4 | 2.03 | 27.1 | 2.41 |
| | Opt-2 | 0.58 | 2.75 | −2.4 | 2.15 | 4.0 | 2.54 | 9.1 | 2.90 |
| | Opt-3 | 0.82 | 4.32 | 22.3 | 4.86 | 13.7 | 3.54 | 8.3 | 3.35 |
| | Opt-4 | 1 | 0 | 0 | 0 | 0 | 0 | 0 | 0 |

Supplementary Tab. 2: Parameters values used for the visiting strategy (see Eqs. 2 and 3 in the main text) for MIMIC agents (collaborator, neutral, and defector), and optimized agents (Opt-1, Opt-2, Opt-3, and Opt-4). These values result from the optimization procedure described in the Materials and Methods section.

| | Col | Neu | Def | |
|---|---|---|---|---|
| $\overline{\text{Col}}$ | 96 % | 4 % | 0 % | 84 % |
| $\overline{\text{Neu}}$ | 9 % | 72 % | 8 % | 13 % |
| $\overline{\text{Def}}$ | 0 % | 21 % | 79 % | 3 % |
| | 84 % | 13 % | 3 % | |

(A) Rule 1

| | Col | Neu | Def | |
|---|---|---|---|---|
| $\overline{\text{Col}}$ | 70 % | 28 % | 1 % | 11 % |
| $\overline{\text{Neu}}$ | 9 % | 69 % | 22 % | 61 % |
| $\overline{\text{Def}}$ | 1 % | 11 % | 88 % | 28 % |
| | 13 % | 49 % | 38 % | |

(B) Rule 2

Supplementary Tab. 3: Fractions of behavioral profiles adopted by participants, whether it is calculated on a single experimental run or over the ten experimental runs (average behavioral profile). In the table, col, neu, and def correspond respectively to collaborators, neutrals, and defectors. The lines above col, neu, and def indicate the average profiles.
Observing the table row-wise reveals that individuals tend to maintain a consistent behavioral profile across the ten experiments. For instance, in Rule 2, an individual who has adopted on average a collaborator profile across the ten experiments was respectively a collaborator 70 % of the experiments, a neutral 9 % of the experiments, and a defector 1 % of the experiments. By examining only the total fractions, shown in the bottom row and right column, one can observe that for each behavioral profile, these fractions remain the same whether they are calculated in single experimental runs or across the ten experiments in Rule 1, and quite similar in Rule 2.

### Supplementary Movies

These movies can be downloaded from the following link:
https://www.dropbox.com/sh/d09oaj0omug4hha/AADFDVE7ycF1RsTmgC6LwRlQa?dl=0.

**Supplementary Movie 1:** Dynamics of ratings $P(t)$ (in red) and visits $Q(t)$ (in blue), as a function of the round $t$, for Rule 1. $(A)$ and $(D)$ The first column corresponds to an experiment where the group of 5 participants achieved the final normalized score $\hat{S}(t = 20)/\hat{S}_{\max} \approx 0.24$ (where $\hat{S}_{\max} = 5420 \times 5 = 27100$ is the maximum possible group score). $(B)$ and $(E)$ The second column corresponds to a simulation of the model where a group of 5 MIMIC agents also obtained a normalized score close to 0.24. Note: $(A)$ corresponds exactly to what the actual participant saw during the experiment.

**Supplementary Movie 2:** Dynamics of ratings $P(t)$ (in red) and visits $Q(t)$ (in blue), as a function of the round $t$, for Rule 2. $(A)$ and $(D)$ The first column corresponds to an experiment where the group of 5 participants achieved the final normalized score $\hat{S}(t = 20)/\hat{S}_{\max} \approx 0.40$ (where $\hat{S}_{\max} = 5420 \times 5 = 27100$ is the maximum possible group score). $(B)$ and $(E)$ The second column corresponds to a simulation of the model where a group of 5 MIMIC agents also obtained a normalized score close to 0.40. Note: $(A)$ corresponds exactly to what the actual participant saw during the experiment.

### Supplementary Data

**Supplementary Data 1:** All data needed to evaluate and replicate the conclusions of the article are present in the article, the Supplementary Materials, or available at the following online repository: https://github.com/Thomas-bssnt/Stigmer-article.git.
